# Autophagy collaborates with apoptosis pathways to control myelination specificity and function

**DOI:** 10.1101/2022.12.31.522394

**Authors:** Tingxin Zhang, Han-Gyu Bae, Aksheev Bhambri, Yihe Zhang, Daniela Barbosa, Jumin Xue, Sabeen Wazir, Sara B. Mulinyawe, Jun Hee Kim, Lu O. Sun

**Affiliations:** Department of Molecular Biology, University of Texas Southwestern Medical Center, Dallas, TX 75390, USA; Graduate Program in Genetics, Development, and Disease; Graduate Program in Neuroscience; Peter O’Donnell Jr. Brain Institute, University of Texas Southwestern Medical Center, Dallas, TX 75390, USA; Hamon Center for Regenerative Science and Medicine, University of Texas Southwestern Medical Center, Dallas, TX 75390, USA; Department of Cellular and Integrative Physiology, University of Texas Health Science Center, San Antonio, TX 78229, USA; Department of Neurology and Neurological Sciences, Stanford University School of Medicine, Stanford, CA 94305, USA

**Keywords:** Oligodendrocyte, myelination, central nervous system, autophagy, apoptosis, compound action potentials, conduction velocity

## Abstract

Oligodendrocytes are the sole myelin producing cells in the central nervous system. Oligodendrocyte numbers are tightly controlled across diverse brain regions to match local axon type and number, but the underlying mechanisms and functional significance remain unclear. Here, we show that autophagy, an evolutionarily conserved cellular process that promotes cell survival under canonical settings, elicits premyelinating oligodendrocyte apoptosis during development and regulates critical aspects of nerve pulse propagation. Autophagy flux is increased in premyelinating oligodendrocytes, and its genetic blockage causes ectopic oligodendrocyte survival throughout the entire brain. Autophagy acts in the TFEB-Bax/Bak pathway and elevates *PUMA* mRNA levels to trigger premyelinating oligodendrocyte apoptosis cell-autonomously. Autophagy continuously functions in the myelinating oligodendrocytes to limit myelin sheath numbers and fine-tune nerve pulse propagation. Our results provide *in vivo* evidence showing that autophagy promotes apoptosis in mammalian cells under physiological conditions and reveal key intrinsic mechanisms governing oligodendrocyte number.

**HIGHLIGHTS:**

- Autophagy flux increases in the premyelinating and myelinating oligodendrocytes
- Autophagy promotes premyelinating oligodendrocyte (pre-OL) apoptosis to control myelination location and timing
- Autophagy acts in the TFEB-PUMA-Bax/Bak pathway and elevates *PUMA* mRNA levels to determine pre-OL fate
- Autophagy continuously functions in the myelinating oligodendrocytes to limit myelin sheath thickness and finetune nerve pulse propagation

## INTRODUCTION

Oligodendrocytes (OLs) are the sole myelin producing cells in the central nervous system (CNS) and are critical for many aspects of neural function (Elbaz and Popko, 2019; Nave and Werner, 2021). Oligodendrocyte numbers are tightly controlled across diverse brain regions to possibly match local axon type and number, however the underlying cellular and molecular mechanisms as well as the functional significance remain elusive. Once oligodendrocyte precursor cells (OPCs) exit the cell cycle, the control of post-mitotic OL number is primarily achieved at an intermediate stage during differentiation, called premyelinating oligodendrocytes (pre-OLs). In rodents, pre-OLs are overproduced and a great number of them undergo programmed cell death before committing to myelination, thereby contributing to spatiotemporal specificity in myelination during development and throughout adulthood (Barres et al., 1992; Sun et al., 2018).

The percentage of pre-OLs that survive and further myelinate axons are brain-region dependent. For instance, about 20-80% pre-OLs in the cortex die early postnatally and throughout adulthood (Hughes et al., 2018; Trapp et al., 1997). In the cerebellar molecular layer, nearly 100% pre-OLs undergo programmed cell death, making this brain region a unique “unmyelinated” area that lacks mature OLs and myelin (Goebbels et al., 2017). Intriguingly, pre-OLs are at a “stressed” differentiation stage with high energy demands; after OPCs differentiate into pre-OLs, they begin expressing large quantities of mRNAs to encode myelin proteins, drastically expand their plasma membrane areas for myelination, and compete against each other for limited nutrients provided by axons and other cell types (Hughes and Stockton, 2021). Previous work showed that the TFEB-PUMA-Bax/Bak pathway powerfully promotes pre-OL apoptosis to control oligodendrocyte number (Sun *et al*., 2018). However, it remains unclear if pre-OL apoptosis is the sole mechanism or other pathways collaborate/counteract with this pathway to determine pre-OL fate.

Macroautophagy (hereafter as autophagy) is an evolutionarily conserved cellular process that degrades unnecessary or dysfunctional cytoplasmic components, thereby promoting cell adaption under nutrient deprivation and stress (Green and Levine, 2014; Griffey and Yamamoto, 2022; Yamamoto and Yue, 2014). Under canonical settings autophagy promotes cell survival, however in rare cases it can elicit cell death directly or indirectly through intimate crosstalk with apoptosis pathways (Denton and Kumar, 2019; Doherty and Baehrecke, 2018). For instance, during *drosophila* midgut development autophagy functions independent of apoptosis to promote midgut cell death, known as autophagy-dependent cell death (Denton et al., 2009). In addition, the selective autophagy of anti-apoptotic proteins, such as dBruce, contributes to the induction of apoptosis during *drosophila* oogenesis (Nezis et al., 2010). In mammals, when apoptosis is genetically blocked autophagy can act as a compensatory mechanism to facilitate elimination of interdigit web cells, thereby contributing to tissue remodeling during early embryogenesis (Arakawa et al., 2017). The *in vivo* evidence showing that autophagy promotes mammalian cell apoptosis under physiologically relevant circumstances, however, is still lacking.

Here, we discovered that autophagy collaborates with apoptosis pathways to eliminate subsets of pre-OLs during development, thereby ensuring spatiotemporal specificity and functional integrity of CNS myelination. We showed that autophagy flux is elevated in the pre-OLs during differentiation, and that genetic perturbation of autophagy within the pre-OLs leads to ectopic myelination in the unmyelinated brain regions, increases oligodendrocyte numbers across diverse brain areas, and alters myelination timing in the forebrain. These phenotypes are due to autophagy’s cell-autonomous function in promoting apoptosis in subsets of pre-OLs. We found that autophagy genetically interacts with the TFEB-PUMA-Bax/Bak pathway and elevates *PUMA/Bbc3* mRNA levels to trigger pre-OL apoptosis. Finally, autophagy continuously functions in the myelinating oligodendrocytes to limit myelin wrap numbers independent of its pro-apoptotic roles in pre-OLs, and it regulates critical aspects of nerve pulse propagation. Our findings showed that autophagy actively promotes pre-OL apoptosis in the developing murine brain, allowing the spatiotemporal specificity and functional integrity of CNS myelination.

## RESULTS

### Autophagy flux is elevated in the premyelinating oligodendrocytes

To determine at which differentiation stage autophagy flux is elevated in oligodendrocyte lineage cells (Figure 1A), we first analyzed the *in vivo* bulk RNA sequencing dataset that contains the transcripts of 399 autophagy genes (gene ontology: 0006919) (Zhang et al., 2014). This dataset covers seven major brain cell types, including the oligodendrocyte precursor cells (OPCs), newly formed oligodendrocyte or premyelinating oligodendrocytes (pre-OLs), and myelinating oligodendrocytes (OLs) (Table S1). We found that autophagy genes exhibited differentiation-stage-dependent expression; many of them started mRNA expression at the OPC and pre-OL stages, whereas a subset of them exhibited the highest mRNA expression in the myelinating OL stage (Figure 1B). To validate the differentiation stage when autophagy flux is elevated, we acutely purified OPCs from early postnatal mouse brains and differentiated them *in vitro* in a serum-free, nutrient-defined culture medium (Figure 1C). The OPCs exhibit stereotypical differentiation in culture; at day 3-4 they display ramified morphologies and gene expression profiles similar to their *in vivo* counterparts known as the pre-OLs, and after day 6 they drastically extend their processes and plasma membrane resembling the myelinating OLs (Figure 1C) (Sun *et al*., 2018). We found that several autophagy proteins, including Lamp2, Beclin1, WIPI2, ATG5, and ATG7, exhibited elevated expressions at differentiation day 4 and remained expressed at day 6 (Figure 1D; quantified in Figure 1F). To address if autophagy flux indeed increases at the pre-OL stage, we treated differentiating OLs with Bafilomycin A1 (BafA1), a potent autophagosome-lysosome fusion blocker that allowed us to analyze transient autophagy flux events (Mauthe et al., 2018); In the presence of BafA1, lysosomal acidification is perturbed which results in the accumulation of autophagosomes that would be degraded during the drug treatment period. We found that in the presence of BafA1 autophagosome adaptor protein p62/SQSTM1 and autophagosome marker LC3-II were significantly increased at the beginning of pre-OL differentiation and throughout its maturation, suggesting that autophagy flux increases at the pre-OL stage (Figure 1E; quantified in Figures 1F and 1G). Finally, we performed transmission electron microscopy (TEM) analysis and found numerous autophagosomes in cultured pre-OLs (Figures 1H-H’; Figure S1; see more examples and quantification in Figures 3 and S3). Therefore, autophagy flux increases during oligodendrocyte differentiation and is elevated in the pre-OLs.

**Figure 1.**
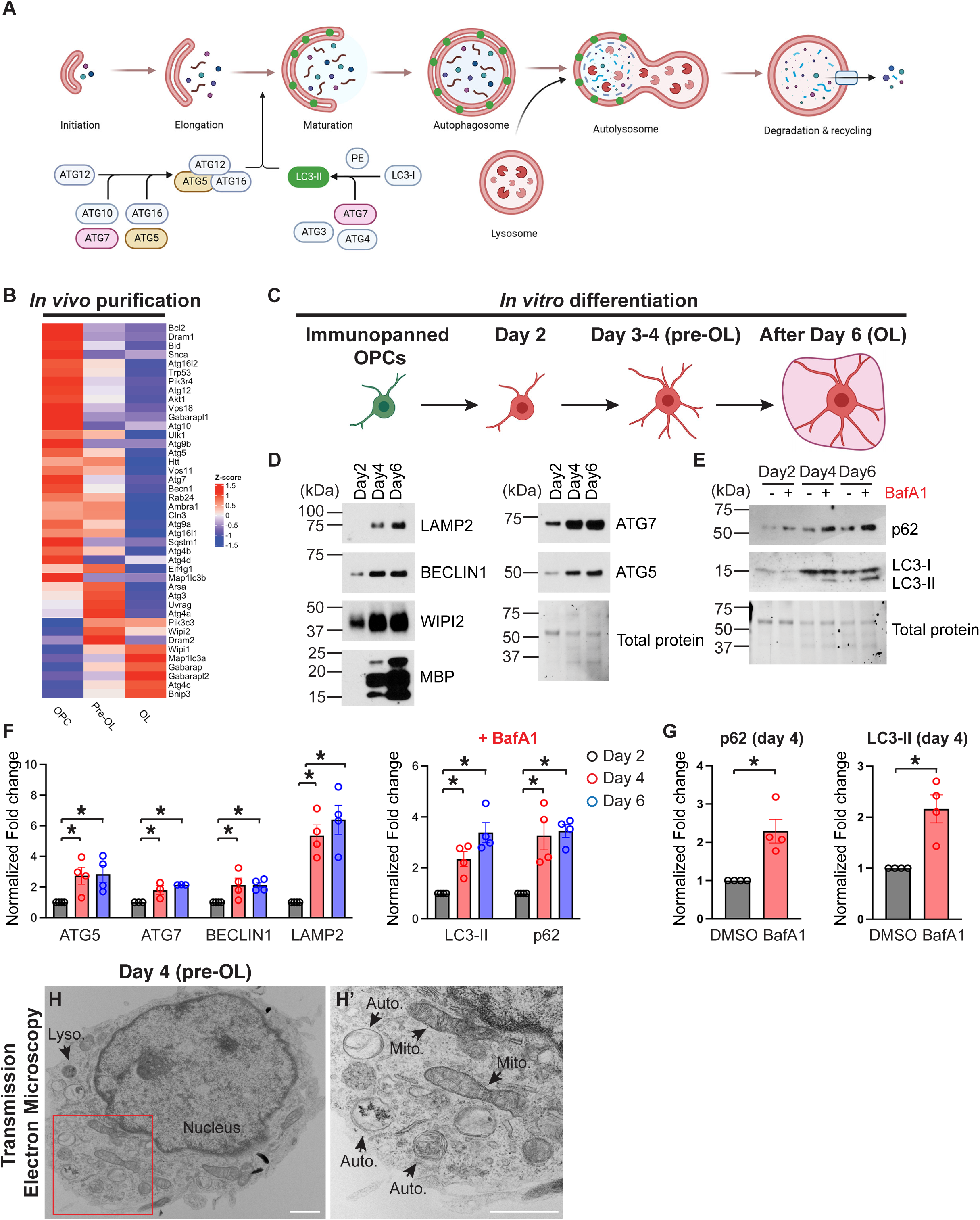
Autophagy flux increases in the premyelinating and myelinating oligodendrocytes. **(A)** Diagram showing the autophagy flux (modified from a template provided by BioRender). ATG5 and ATG7 are the critical mediators for autophagosome elongation. The lipidation of LC3-I followed by the insertion of LC3-II to the autophagosome serve as the marker for autophagy flux. **(B)** Heatmap showing major autophagy gene expression levels (in z scores) in acutely isolated oligodendrocyte precursor cells (OPCs), premyelinating oligodendrocytes (pre-OLs), and myelinating oligodendrocytes (OLs) (Zhang., et al., 2014). **(C)** Diagram of oligodendrocyte *in vitro* differentiation. At differentiation day 3-4, oligodendrocytes exhibit ramified morphologies and gene signatures resembling pre-OLs *in vivo*. After day 6, oligodendrocytes display flattened, “pancake”-like morphologies and similar gene expression profiles as myelinating OLs *in vivo* (Sun *et al*., 2018). **(D-E)** Western blot analysis of autophagy-related protein expression during oligodendrocyte *in vitro* differentiation in the presence or absence of Bafilomycin A1 (BafA1) treatment, showing that autophagy flux increases in pre-OLs and mature OLs. **(F)** Quantification of autophagy protein expression (left panel) and LC3-II and p62 levels in the presence of BafA1 treatment (right panel) during *in vitro* differentiation. Protein levels were normalized to the total protein level. **(G)**. Comparison of p62 (left) and LC3-II (right) between vehicle treatment (DMSO) and BafA1 treatment at differentiation day 4, showing that autophagy flux increases at the pre-OL stage. **(H and H’)** Representative transmission electron microscopy (TEM) micrograph of a pre-OL at differentiation day 4 *in vitro* (H). H’ represents the enlarged view of the red inset in H. Mito., mitochondria. Auto., autophagosome. Lyso., lysosome. See more examples and quantifications in Figures S1, 3 and S3. Error bars represent SEM. Scale bars: 1 μm. \**p*<0.05.

**Figure 2.**
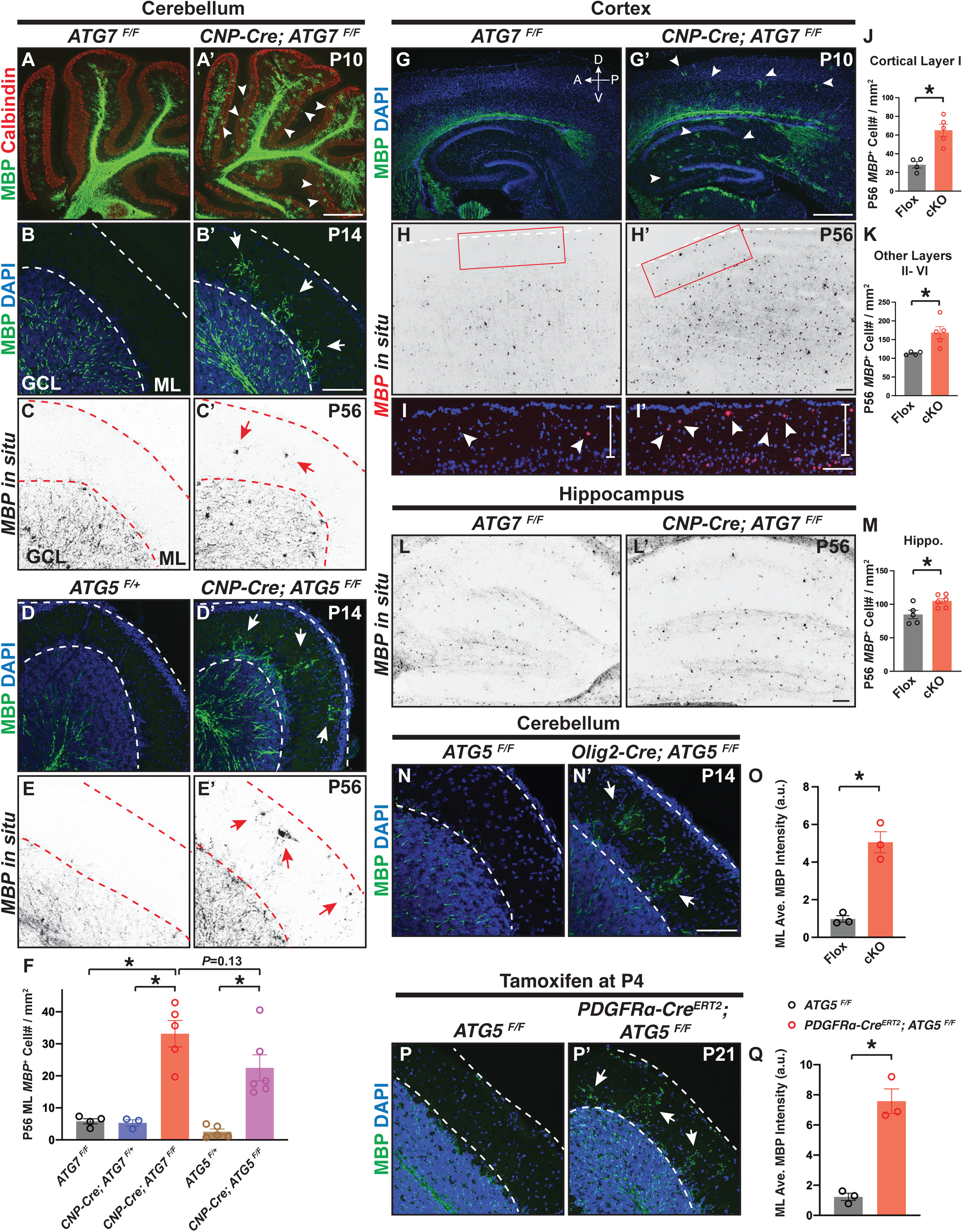
Genetic deletion of *ATG7* or *ATG5* in oligodendrocyte lineage cells disrupts the location and timing of myelination. **(A-B’)** Characterization of ectopic oligodendrocytes and aberrant myelination in the cerebellar molecular layer (ML) in a mutant mouse line where *ATG7* was conditionally deleted in oligodendrocyte lineage cells (*CNP-Cre; ATG7^F/F^*) at P10 (A’) and P14 (B’) as compared to the littermate controls (A and B). White arrowheads in A’ and white arrows in B’ indicate ectopic myelin basic protein (MBP) immunolabeling in the cerebellar ML. GCL, granule cell layer. **(C and C’)** *In situ* hybridization using the probes against *MBP* mRNA showing that *CNP-Cre; ATG7^F/F^* mutants exhibited ectopic *MBP^+^* oligodendrocytes in the cerebellar ML (red arrows in C’). **(D-E’)** Conditional deletion of *ATG5* in oligodendrocyte lineage cells resulted in ectopic MBP immunolabeling (white arrows in D’) and aberrant *MBP^+^* oligodendrocytes (red arrows in E’) in the cerebellar ML at P14 (D and D’) and P56 (E and E’). **(F)** Quantification of *MBP^+^* oligodendrocyte numbers in the cerebellar ML of P56 *ATG7^F/F^*, *CNP-Cre; ATG7^F/+^*, *CNP-Cre; ATG7^F/F^*, *ATG5^F/+^*, and *CNP-Cre; ATG5^F/F^* mice. **(G-I’)** *CNP-Cre; ATG7^F/F^* mutants exhibited MBP immunolabeling precociously in the cortex and hippocampus at P10 (white arrowheads in G’; see also Figure S2H’; n=3 animals per genotype). At P56, *CNP-Cre; ATG7^F/F^* mutants displayed increased *MBP^+^* oligodendrocytes in cortical layer I (red inset in H’ and white arrowheads in I’) as compared to littermate controls (red inset in H and white arrowheads in I). I and I’ represent the insets shown in H and H’, respectively. A, anterior; P, posterior; D, dorsal; V, ventral. **(J and K)** Quantification of *MBP^+^* oligodendrocytes in cortical layer I (J) and cortical layer II-VI (K) of *ATG7^F/F^* (Flox) and *CNP-Cre; ATG7^F/F^* (cKO) mice at P56. **(L-M)** Representative images (L and L’) and quantification (M) showing that P56 *CNP-Cre; ATG7^F/F^* mutants exhibited significantly increased *MBP^+^* oligodendrocyte numbers in the hippocampus as compared to littermate controls. **(N-O)** Representative images (N and N’) and quantification (O) showing that *Olig2-Cre* mediated deletion of *ATG5* in oligodendrocyte lineage cells resulted in ectopic MBP immunolabeling in the cerebellar ML. **(P-Q)** Representative confocal micrographs (P and P’) and quantification (Q) showing that inducible deletion of *ATG5* from oligodendrocyte precursor cells at P4 led to ectopic MBP immunolabeling in P21 cerebellar ML. Error bars represent SEM. Scale bars: 500 μm in (A’) for (A) and (A’); 100 μm in (B’) for (B)-(E’); 500 μm in (G’) for (G)-(G’); 100 μm in (H’) for (H)-(H’); 100 μm in (I’) for (I)-(I’); 100 μm in (L’) for (L)-(L’); and 100 μm in (N’) for (N), (N’), (P), and (P’). \**p*<0.05.

**Figure 3.**
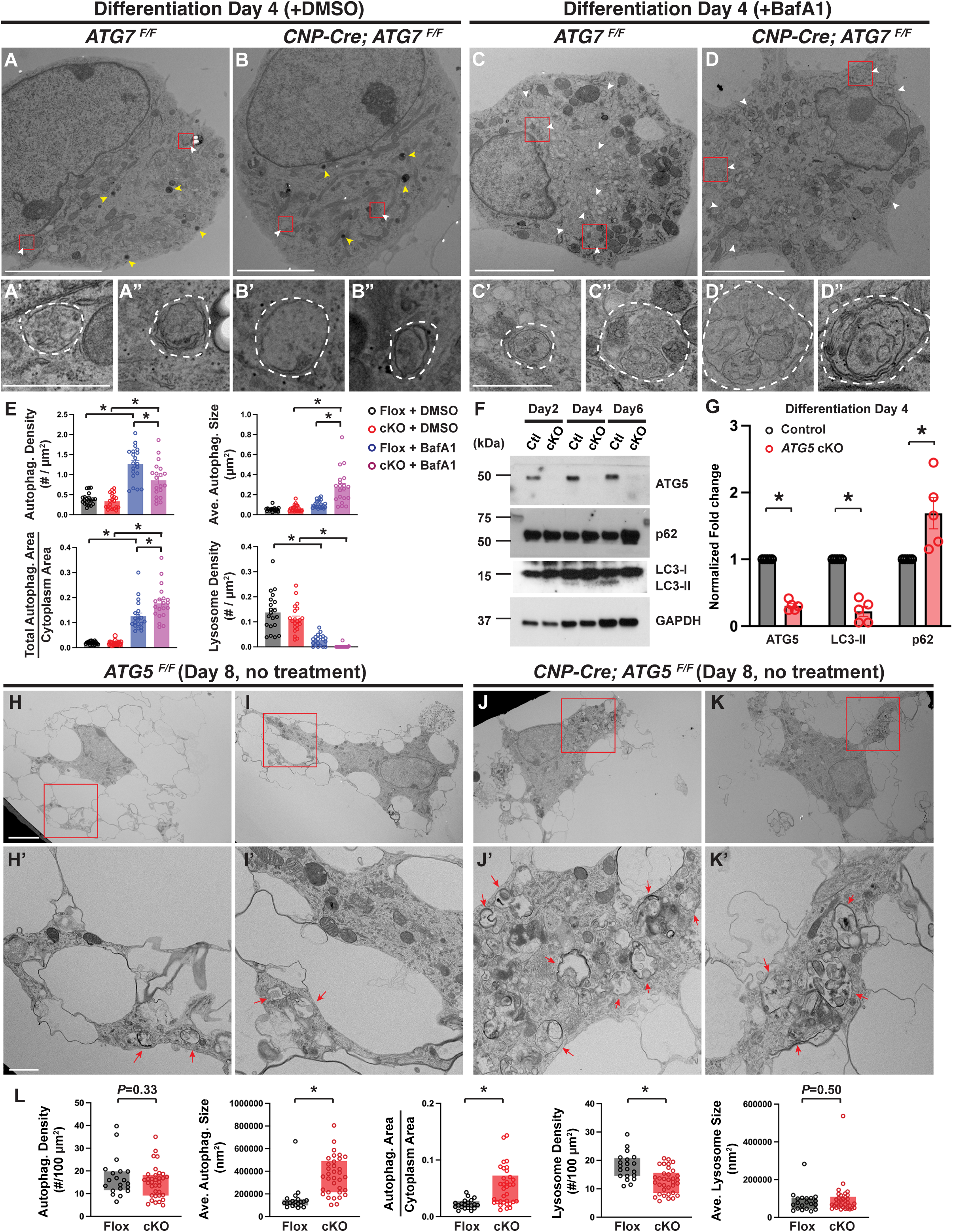
*ATG5*- and *ATG7*-deficient premyelinating oligodendrocytes exhibit disrupted autophagy flux and autophagosome abnormalities. **(A-D”)** Transmission electron microscopy (TEM) analysis of *ATG7^F/F^* oligodendrocytes (A-A” and C-C”) and *CNP-Cre; ATG7^F/F^* oligodendrocytes (B-B” and D-D”) in the presence of dimethyl sulfoxide (DMSO; A-B”) or DMSO containing 50nM Bafilomycin A1 (BafA1; C-D”) at differentiation day 4. A’-D” represent the red insets shown in A-D, respectively. Yellow arrowheads indicate lysosomes. White arrowheads indicate autophagosomes. Dashed lines in A’-D” demarcate autophagosomes. **(E)** Quantifications of autophagosome density (top left), average autophagosome size (top right), ratio of total autophagosome area over cytoplasm area (bottom left), and lysosome density (bottom right) in oligodendrocytes at differentiation day 4 when they were at the pre-myelinating stage *in vitro*. n≥20 cells per condition per genotype. **(F and G)** Western blot analysis (F) and quantification (G) of ATG5, autophagy adaptor protein p62, and LC3-II in *ATG5^F/F^* (control) and *CNP-Cre; ATG5^F/F^* (*ATG5* cKO) oligodendrocytes during *in vitro* differentiation, showing that LC3-II levels were significantly reduced whereas p62 levels were increased in the absence of ATG5. Protein levels were normalized to GAPDH. **(H-K’)** Representative TEM images of *ATG5^F/F^* oligodendrocytes (H, I, H’, and I’) and *CNP-Cre; ATG5^F/F^* oligodendrocytes (J, K, J’, and K’) at differentiation day 8, showing that autophagosomes exhibited defective morphologies and enlarged sizes in *CNP-Cre; ATG5^F/F^* oligodendrocytes (red arrows in J’ and K’). H’-K’ represent red insets in H-K, respectively. **(L)** Quantifications (box and whisker, min to max) of autophagosome density, average autophagosome size, the ratio of total autophagosome area over cytoplasm area, lysosome density, and average lysosome size in *ATG5^F/F^* (Flox) and *CNP-Cre; ATG5^F/F^* (cKO) oligodendrocytes at differentiation day 8. n≥20 cells per condition per genotype. Error bars represent SEM. Scale bars: 5 μm in (A)-(D); 1 μm in (A’) for (A’)-(B”); 1 μm in (C’) for (C’)- (D”); 5 μm in (H) for (H)-(K); and 1 μm in (H’) for (H’)-(K’). \**p*<0.05.

### Genetic blockade of autophagy flux in oligodendrocytes disrupts myelination location and timing

To determine the roles of autophagy in pre-OLs, we characterized mouse cerebellum, a brain region that harbors numerous pre-OLs during development (Sun *et al*., 2018). The cerebellum white matter tracts are heavily myelinated, however the cerebellar molecular layer (ML) is not myelinated despite the fact that a great number of cerebellar granule cell axons extend throughout the ML (Goebbels *et al*., 2017). Intriguingly, OPCs exist in the cerebellar ML and can differentiate into pre-OLs, however none of these pre-OLs survive in the ML, making the cerebellar ML an un-myelinated brain region (Sun *et al*., 2018). To determine if autophagy regulates pre-OL viability in the cerebellar ML, we conditionally deleted *ATG7*, a critical gene mediating autophagosome elongation, in oligodendrocyte lineage cells (*CNP-Cre; ATG7^F/F^*). Indeed, myelin basic protein (MBP) as well as myelination were restricted beneath the cerebellar ML in littermate control animals (Figures 2A-B, Figure S2G). In strong contrast, *CNP-Cre; ATG7^F/F^* mutants exhibited fully-penetrant, ectopic presence of myelin proteins and myelin wrapping in the ML across the entire cerebellum (Figures 2A’-B’, Figures S2G’-G”). To determine if the aberrant myelin was caused by ectopic OLs in the ML, we utilized *in situ* hybridization with probes recognizing *MBP* mRNAs, a marker expressed by premyelinating and myelinating OLs (Xiao et al., 2016). We found that at postnatal day 56 (P56) both *ATG7^F/F^* and *CNP-Cre; ATG7^F/+^* mice harbored very few *MBP^+^* OLs in the cerebellar ML, likely representing newly formed pre-OLs that haven’t yet undergone programmed cell death (Figure 2C; quantified in Figure 2F). Conversely, *ATG7* cKO mice exhibited significantly increased numbers of *MBP^+^* cells in the ML (Figure 2C’; quantified in Figure 2F; 5.8±0.8 cells/mm^2^ in *ATG7^F/F^* ML, 5.3±0.9 in *CNP-Cre; ATG7^F/+^*, and 33.1±4.1 cells/mm^2^ in *ATG7* cKO, mean±SEM, n≥3 animals per genotype, \**p*<0.001, one-way ANOVA followed with Tukey’s multiple comparisons test). We found similar phenotypes in the *CNP-Cre; ATG5^F/F^* mutants (Figures 2D-E’; quantified in 2F) and *Olig2-Cre; ATG5^F/F^* mutants (Figures 2N-N’; quantified in Figure 2O). Because *CNP-Cre* and *Olig2-Cre* are transiently expressed in neurons (Tognatta et al., 2017; Zawadzka et al., 2010), the ectopic myelination phenotype observed in the *CNP-Cre; ATG5^F/F^, CNP-Cre; ATG7^F/F^*, or *Olig2-Cre; ATG5^F/F^* mutants could be due to neuronal disruption of autophagy (Jo et al., 2021). To rule out this possibility, we conditionally deleted *ATG5* only in OPCs from P4 by administering tamoxifen to *PDGFRα-Cre^ERT2^; ATG5^F/F^* mice. We found identical phenotypes in these mutants (Figures 2P-P’; quantified in Figure 2Q), together showing that the ectopic myelination phenotype is indeed due to autophagy disruption within oligodendrocyte lineage cells.

Pre-OLs are widespread in the brain during myelination and serve as a critical transition stage between OPCs and myelinating OLs (Hughes and Stockton, 2021). In the cortex, pre-OLs are overproduced and nearly 80% of them are eliminated by apoptosis (Hughes *et al*., 2018). To determine if autophagy functions globally in the pre-OLs to control oligodendrocyte number, we characterized a variety of brain regions including the cortex and hippocampus throughout development. Myelination in the wild-type forebrain didn’t start until P7, with only a few brain areas harboring differentiating MBP^+^ OLs (Figure S2H and Figure 2G). In strong contrast, *ATG7* cKO mice exhibited a fully-penetrant, “precocious” myelination phenotype where numerous MBP^+^ OLs were found throughout the entire forebrain as early as P7 (Figure S2H’). By P10 MBP^+^ OLs readily appeared in the deep cortical layers and hippocampus in *ATG7* cKO mice, whereas the littermate controls didn’t contain any MBP^+^ OLs in those brain regions (Figure 2G’). This phenotype lasted until adulthood, as both P56 *ATG7* cKO and *ATG5* cKO mice exhibited significantly increased myelinating OL numbers in the cortex and hippocampus (Figures 2H-M, S2A-C, and S2F-F’). Importantly, these phenotypes were not due to dysregulation of OPC migration, patterning, or proliferation, because OPC distribution, number, as well as proliferation rate were not changed in *ATG7* cKO mice (Figures S2D-E and S2I-K). Thus, genetic perturbation of autophagy flux in oligodendrocyte lineage cells generates ectopic OLs during development and into adulthood, resulting in disrupted myelination location and timing.

### Autophagy functions cell-autonomously to promote apoptosis in subsets of premyelinating oligodendrocytes

What are the cellular mechanisms underlying autophagy-dependent elimination of pre-OLs? We began to address this question by analyzing autophagy flux in differentiating oligodendrocytes with biochemical analysis and high-resolution TEM. Consistent with the reported role of ATG5 in mediating autophagy flux, we found that autophagy adaptor protein p62 were significantly increased and that autophagosome marker LC3-II displayed a significant decrease at differentiation day 4 (pre-OL stage) and day 8 (mature OL stage) in *ATG5* cKO OLs as compared to control cells (Figure 3F; quantified in Figure 3G). In addition, TEM analysis revealed that *ATG5* cKO pre-OLs exhibited a significantly reduced lysosomal density (Figures S3A-B”; quantified in Figure S3E), showing that autophagy flux was disrupted in *ATG5* cKO pre-OLs.

Because autophagosomes undergo continuous maturation and fusion with lysosomes (Klionsky et al., 2021), snapshots of the fixed OLs may not reveal detailed abnormalities in autophagy flux (Figures S3C-D). To determine if pre-OL autophagosomes are dysregulated in the absence of ATG5 or ATG7, we applied BafA1 briefly in culture to temporarily inhibit autophagy flux in the pre-OLs. Indeed, autophagosome numbers were significantly increased and lysosomes were abolished in BafA1-treated cells, showing that BafA1 was effective in blocking autophagy flux (compare Figures 3A-B with Figures 3C-D; quantified in Figure 3E). We found that in the presence of BafA1 *ATG7* deficient pre-OLs (*CNP-Cre; ATG7^F/F^*) displayed significantly reduced autophagosome densities and increased autophagosome sizes (quantified in Figure 3E and Figure S3G). Notably, in the presence of BafA1 *ATG7* cKO pre-OLs harbored increased numbers of large degradative compartments (DGCs) that contained autophagosomes, amphisomes, and autolysosomes (compare Figures 3C’-C” with Figures 3D’-D”; quantified in Figure 3E). To rule out the possibility that these ultrastructural deficits were due to BafA1 treatment (Mauthe *et al*., 2018), we further differentiated oligodendrocytes until day 8 when autophagy flux deficits had manifested over a long period of time in culture. We found that without any treatment *ATG5* cKO OLs readily exhibited similar autophagosome abnormalities as observed at differentiation day 4 (Figures 3H-K’ and Figures S3H-O). These deficits include significantly increased autophagosome sizes and the total area, as well as reduced lysosome densities (Figure 3L). These observations were in agreement with the canonical functions of ATG5 and ATG7 where they mediate the elongation of autophagosomes (Kishi-Itakura et al., 2014); in the absence of ATG5/7, autophagosomes may continue enclosing cellular contents and organelles without proper closures, leading to increased autophagosome areas in the pre-OLs. Thus, *ATG5-* and *ATG7*-deficient OLs exhibited disrupted autophagy flux and aberrant autophagosomes during differentiation.

Under canonical settings, autophagy promotes cell adaption and survival upon stress and starvation (Griffey and Yamamoto, 2022). Our data showed that, however, *ATG5* and *ATG7* cKO mice exhibited ectopic OLs in the non-myelinated cerebellar molecular layer (ML) and aberrantly increased OL numbers in other brain regions (Figure 2). These results suggest that autophagy promotes OL cell death at the pre-OL stage. To test this hypothesis, we first analyzed the cerebellar ML at P11 when wild-type pre-OLs were transiently differentiated and rapidly underwent programmed cell death (Figures 4B) (Sun *et al*., 2018). By using an antibody raised against MBP which begins its expression in the pre-OLs (Xiao *et al*., 2016), we found that a fraction of MBP^+^ pre-OLs were cleaved caspase3^+^ in *ATG5^F/F^* cerebellar ML (Figures 4B’-B”; quantified in Figure 4E). In contrast, although MBP^+^ cleaved caspase-3^+^ double positive cells were occasionally found in *ATG5* cKO cerebellar ML (Figures 4D-D”), the ratio of MBP^+^ and cleaved-caspase 3^+^ double positive cells over total MBP^+^ cells in *ATG5* cKO mice was significantly reduced as compared to littermate controls (Figures 4C-C”; quantified in Figure 4E; ratio=8.2±2.3% for *ATG5^F/F^* mice and 0.8±0.4% for *CNP-Cre; ATG5^F/F^* mice, Mean±SEM, n=3 animals per genotype, \**p*=0.03 by two-tailed unpaired *t*-test). We found a similar reduction in P10 *ATG7* cKO mice (quantified in Figure 4F; 7.8±2.5% for *ATG7^F/F^* and 2.4±1.5% for *CNP-Cre; ATG7^F/F^*, Mean±SEM, n=4 animals per genotype, \**p*=0.01 by two-tailed unpaired *t*-test), together showing that pre-OL apoptosis was reduced in the cerebellar ML when autophagy flux was genetically perturbed in these cells.

**Figure 4.**
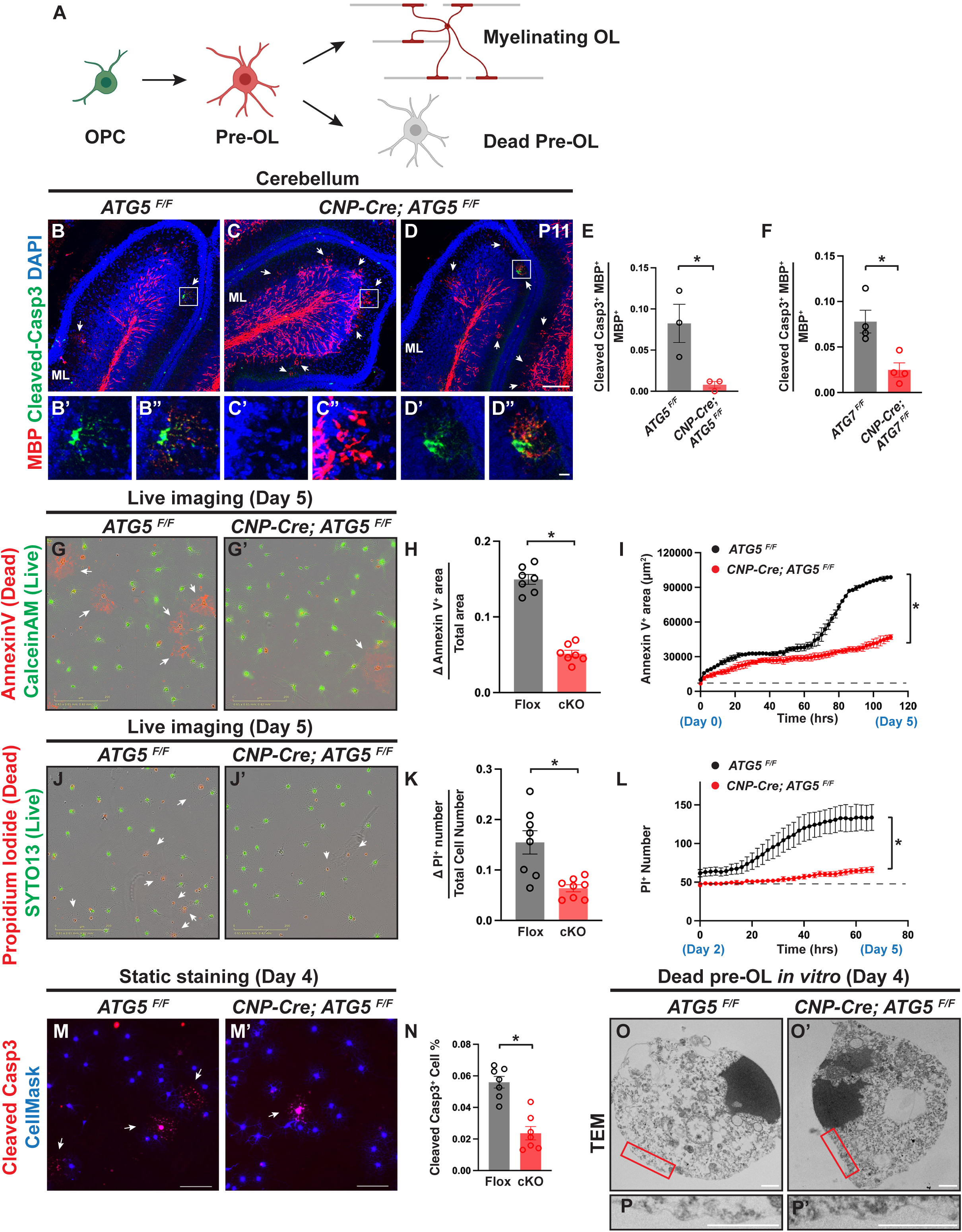
Autophagy functions cell-autonomously to promote premyelinating oligodendrocyte apoptosis. **(A)** Diagram showing that a great fraction of oligodendrocytes undergoes programmed cell death at the premyelinating stage before committing to myelination. **(B-D and B’-D”)** Representative confocal micrographs of P11 cerebellums from *ATG5^F/F^* (B-B”) and *CNP-Cre; ATG5^F/F^* (C-D”) showing that a fraction of MBP^+^ pre-OLs in *ATG5^F/F^* cerebellar ML were cleaved caspase3^+^, suggesting that they were undergoing apoptosis. In contrast, most MBP^+^ pre-OLs in *CNP-Cre; ATG5^F/F^* cerebellar ML were cleaved caspase 3^−^ (C-C”) and had proceeded to differentiation in the ML (see also Figures 2B’ and 2C’), whereas very few MBP^+^ cells were cleaved caspase3^+^ (D-D”). B’-B”, C’-C”, and D’-D” represent the insets in B, C, and D, respectively. White arrows indicate transiently differentiated pre-OLs in the cerebellar ML at P11. **(E and F)** Quantification of the ratio of cleaved caspase3^+^ and MBP^+^ double immunopositive cells over total MBP^+^ cells in the cerebellar ML from P11 *ATG5* cKO (E) and P10 *ATG7* cKO (F) mice. (**G and G’**) Representative live-cell imaging micrographs of *ATG5^F/F^* (G) and *CNP-Cre; ATG5^F/F^* oligodendrocytes (G’) at differentiation day 5, showing that pre-OL cell death was reduced in the absence of ATG5. White arrows indicate dead oligodendrocytes delineated by Alexa-555 conjugated Annexin V (red). **(H)** Quantification of the increased annexin V^+^ area from day 3 to day 5 (ΔAnnexin V^+^ area, reflecting pre-OL cell death) over total cell area at differentiation day 5. **(I)** Representative curves of Annexin V^+^ total area throughout the entire live-cell imaging course (from day 0 to day 5), showing that control OLs (black) exhibited drastically increased cell death events beginning at day 3 (∼70 hrs) when the OLs differentiated into the premyelinating stage. In contrast, *CNP-Cre; ATG5^F/F^* mutant cells exhibited significantly reduced cell death (red). \**p*<0.0001 by paired two-tailed *t*-test. The experiment was repeated at least four times with similar results. **(J and J’)** Representative live-cell imaging micrographs of *ATG5^F/F^* (J) and *CNP-Cre; ATG5^F/F^* (J’) at differentiation day 5 labeled by propidium iodide (PI, red for dead cells) and SYTO13 (green for live cells). **(K)** Quantification of the increased PI^+^ cell number from day 2 to day 5 (ΔPI^+^ number, reflecting pre-OL cell death) to the total cell number at day 5 at differentiation day 5. **(L)** Representative curves of PI^+^ cell number from differentiation day 2 (before pre-OL stage) to day 5 (after pre-OL stage), showing that *CNP-Cre; ATG5^F/F^* mutant pre-OLs (red) exhibited significantly reduced cell death as compared to control cells (black), similarly to the *in vivo* observations (see Figures 4C-D”). \**p*<0.0001 by paired two-tailed *t*-test. The experiment was repeated at least three times with similar results. **(M and M’)** Representative micrographs of *ATG5^F/F^* (M) and *CNP-Cre; ATG5^F/F^* oligodendrocytes (M’) immunostained by an antibody raised against cleaved caspase-3 (red) and counterstained by CellMask (blue) at differentiation day 4. **(N)** Quantification of cleaved caspase-3^+^ cell number over total cell number at differentiation day 4. **(O-P’)** Representative TEM micrographs of *ATG5^F/F^* (O and P) and *CNP-Cre; ATG5^F/F^* (O’ and P’) oligodendrocytes at differentiation day 4. The dead cells from both genotypes exhibited nucleus condensation and intact membranes, which are the apoptosis characteristics. P and P’ represent the red insets in O and O’, respectively. Error bars represent SEM. Scale bars: 100 μm in (D) for (B)-(D); 10 μm in (D”) for (B’)-(D”); 200 μm in (G), (G’), (J), and (J’); 100 μm in (M) and (M’); and 1 μm in (O)-(P’). \**p*<0.05.

To determine if autophagy functions cell-autonomously to promote pre-OL apoptosis, we acutely purified OPCs from *ATG5* cKO and littermate control’s brains and differentiated them in a serum-free, nutrient-defined culture medium. Importantly, this culture only contained OPCs but not neurons or other cell types, allowing us to define the autonomous function of autophagy in oligodendrocytes. We performed live-cell imaging assays from the beginning of OPC differentiation (day 0) to day 5 when the cells fully differentiated to OLs. We utilized a fluorescent dye-conjugated Annexin V probe (Alexa-594 Annexin V) to monitor apoptosis during OL differentiation (Sun *et al*., 2018). As expected, we found that a subset of control OLs were Annexin V^+^ on day 3 when they first differentiated into the pre-OL stage (Figure 4I; see also Movie S1). In strong contrast, *ATG5* cKO OLs exhibited significantly reduced Annexin V^+^ cell areas (compare Figures 4G and 4G’; quantified in Figures 4H and 4I; see also Movie S2). This result was further confirmed by the live-cell imaging assay using another cell death marker, propidium iodide (PI), which labels dead cell nuclei (Figures 4J and 4J’; quantified in Figures 4K and 4L), as well as by immunostaining of cleaved caspase 3 in the pre-OLs (Figures 4M and 4M’; quantified in Figure 4N). Importantly, the reduction of apoptosis in *ATG5* cKO pre-OLs was not due to accelerated OPC->pre-OL differentiation, as the percentages of MBP^+^ and galactocerebroside^+^ (GalC^+^) immunopositive cells were not changed in *ATG5* cKO OLs (Figure S4A-F). Finally, to rule out the possibility that pre-OLs undergo a different form of programmed cell death other than apoptosis, we analyzed dead pre-OL’s ultrastructure by TEM. Indeed, wild-type dead pre-OLs exhibited apoptotic characteristics including chromatin condensation and maintenance of intact plasma membrane (Figures 4O and 4P; Figures S4G-J). Similarly, unlike autosis which exhibits swollen perinuclear space or necrosis where plasma membrane erupts (Koenig et al., 2020; Liu et al., 2013), we found that dead *ATG5* cKO pre-OLs still displayed apoptotic characteristics (Figures 4O’ and 4P’; Figures S4G’-J’). Taken together, our results showed that autophagy functions cell-autonomously to promote apoptosis in subsets of pre-OLs.

### Autophagy acts in the TFEB-Bax/Bak pathway to promote pre-OL apoptosis by elevating *PUMA* mRNA levels

To determine the molecular mechanisms through which autophagy promotes pre-OL apoptosis, we attempted to identify proteins that were mis-regulated when autophagy flux was disrupted in the pre-OLs. Because autophagy eliminates unnecessary/dysfunctional proteins and organelles through lysosomal degradation, we employed quantitative liquid chromatography with tandem mass spectrometry (LC-MS-MS) to detect proteins that were aberrantly enriched in *ATG5* cKO pre-OLs (Figure 5A). Indeed, ATG5 was among the most downregulated proteins and that autophagy adaptor protein p62 and Gabarapl2/ATG8 were significantly upregulated in *ATG5* cKO pre-OLs, showing that the assay was successful (Figure 5B). We found that proteins involved in ERphagy (Calcoco1and Retreg1), protein kinase A activity (Prkar1a), and lipid biosynthesis and metabolism were significantly upregulated in *ATG5* cKO OLs (Figures 5B and S6A; Table S2). However, this assay didn’t provide clear indication of the candidates directly involved in the apoptosis pathways, likely due to its limited sensitivity. To address the mechanisms underlying autophagy-mediated pre-OL apoptosis, we employed bulk RNA sequencing (RNA-seq) approach to quantitatively measure transcriptomic changes when autophagy flux was perturbed (Table S3). We focused on apoptosis pathways and found that genes directly involved in apoptosis, including *Pmaip1* (*Noxa*) and *PUMA* (*Bbc3*), exhibited significant decreased expressions in *ATG5* cKO pre-OLs (Figure 5C). We further validated that mRNA and protein levels of *PUMA*, a potent pro-apoptotic gene belonging to the Bcl-2 family, were indeed significantly reduced in *ATG5* cKO pre-OLs (Figures 5D and 5E). Consistent with the RNA sequencing results, major pro- and anti-apoptotic proteins including Bax, Bak, Bcl2, and XIAP remained unchanged in *ATG5* cKO pre-OLs (Figures 5D and 5F).

**Figure 5.**
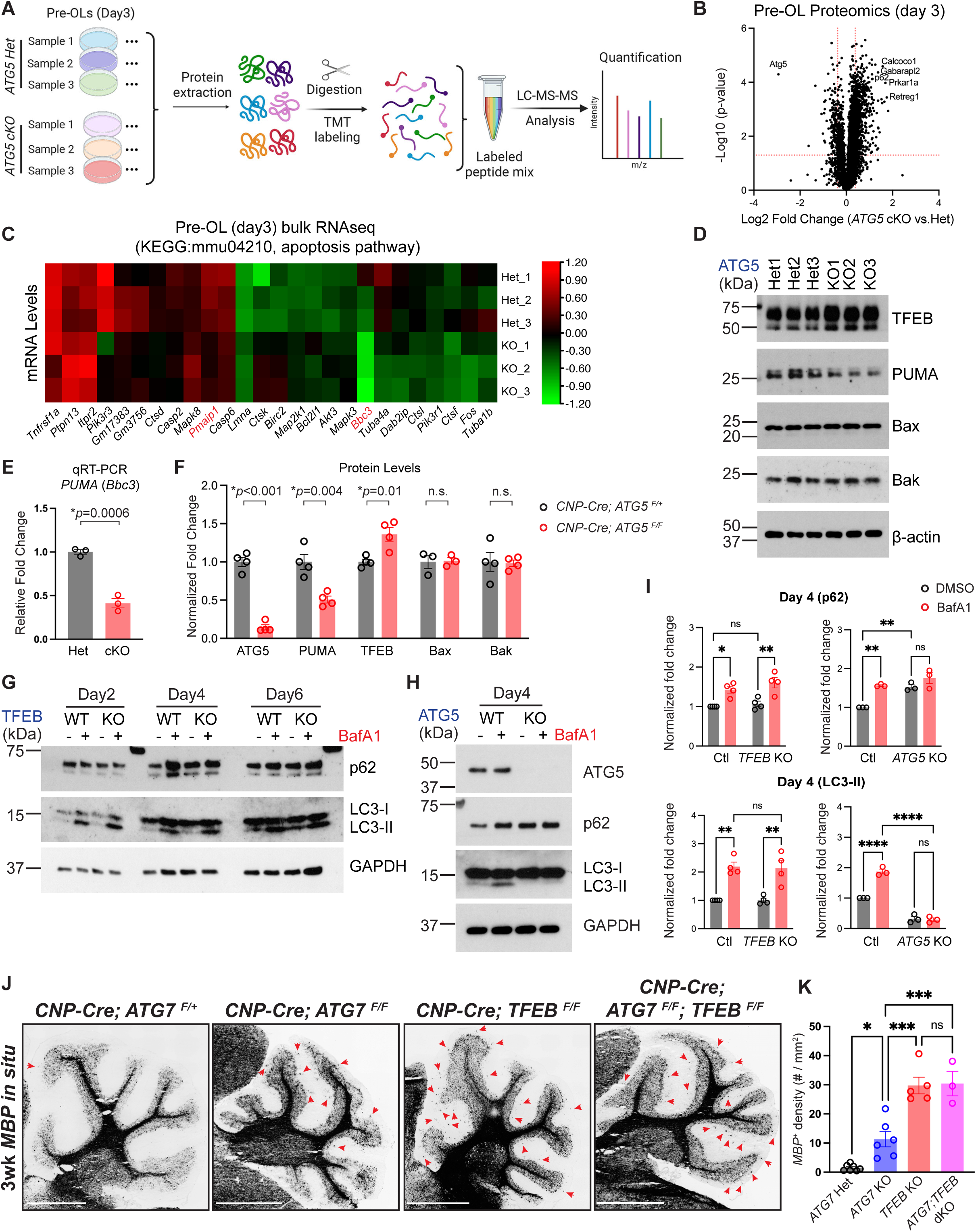
Autophagy acts in the TFEB-PUMA-Bax/Bak pathway and elevates *PUMA* mRNA levels to trigger pre-OL apoptosis. **(A)** Schematics of quantitative liquid chromatography with tandem mass spectrometry (LC-MS-MS) to identify candidates involved in autophagy-mediated pre-OL apoptosis. **(B)** Volcano plot showing significantly enriched and depleted proteins in *CNP-Cre; ATG5^F/F^* (*ATG5* cKO) pre-OLs as compared to the ones in *CNP-Cre; ATG5^F/+^* (*ATG5* Het) at differentiation day 3. Each dot represents a protein. See Table S2 for the full protein list. **(C)** Heatmap showing expression levels (in z scores) of genes belonging to the KEGG mmu04210: apoptosis pathway that were significantly up- or down-regulated (adjusted *p* value<0.05) in *ATG5* cKO pre-OLs at differentiation day 3 by RNA sequencing (RNAseq). Each row represents a biological repeat, and each column represents a gene. *Pmaip1* (*NOXA*) and *Bbc3* (*PUMA*) were the two genes directly associated with apoptosis that exhibited significantly decreased expression in *ATG5* KO pre-OLs. **(D)** Western blot analysis of TFEB, PUMA, Bax, and Bak protein levels in *ATG5* Het and *ATG5* cKO pre-OLs at differentiation day 3, showing that PUMA protein levels were significantly reduced in *ATG5* cKO pre-OLs. See quantification in F. **(E)** Quantitative RT-PCR showing that *PUMA* mRNA levels were significantly reduced in *ATG5* cKO pre-OLs as compared to control cells. **(F)** Quantification of ATG5, TFEB, PUMA, Bax, and Bak protein levels in *ATG5* Het and *ATG5* cKO pre-OLs at differentiation day 3. **(G)** Biochemical analysis of autophagy flux in *TFEB* KO oligodendrocytes in the presence or absence of BafA1, showing that *TFEB* KO oligodendrocytes still retained similar levels of p62 and LC3-II as compared to WT cells under both conditions. See quantification in I. **(H)** Western blot analysis of p62 and LC3-I/II levels in the presence or absence of BafA1 in *ATG5* WT (*ATG5^F/F^*) and KO (*CNP-Cre; ATG5^F/F^*) pre-OLs. As expected, p62 levels were significantly increased in *ATG5* KO pre-OLs as compared to WT cells, whereas LC3-II was abolished in the presence or absence of BafA1 in *ATG5* KO pre-OLs, together showing that autophagy flux was disrupted in *ATG5* KO pre-OLs. See quantification in I. **(I)** Quantification of p62 and LC3-II protein levels in *TFEB* control (*TFEB^F/F^*) *TFEB* KO (*Olig2-Cre; TFEB^F/F^*), *ATG5* control (*ATG5^F/F^*), and *ATG5* KO (*CNP-Cre; ATG5^F/F^*) oligodendrocytes at differentiation day 4 (pre-OL stage) in the presence or absence of BafA1. The protein levels were normalized to GAPDH. Two-way ANOVA followed by Sidak’s multiple comparisons test. **(J)** Representative micrographs from 3-week-old *CNP-Cre; ATG7^F/+^* (*ATG7* Het), *CNP-Cre; ATG7^F/F^* (*ATG7* KO), *CNP-Cre; TFEB^F/F^* (*TFEB* KO), and *CNP-Cre; ATG7^F/F^; TFEB^F/F^* (*ATG7; TFEB* dKO) cerebellums that were labeled by *MBP in situ* probes. Red arrows indicate ectopic oligodendrocytes in the cerebellar ML. **(K)** Quantification of *MBP^+^* oligodendrocyte density in the cerebellar ML, showing that both *TFEB* KO and *ATG7; TFEB* dKO mice exhibited significantly increased oligodendrocyte numbers as compared to *ATG7* KO mice, and that *TFEB* KO and *ATG7; TFEB* dKO mice were indistinguishable. One-way ANOVA followed by Tukey’s multiple comparisons test. Error bars represent SEM. Scale bars: 1 mm in (J). \**p*<0.05, \*\**p*<0.01, \*\*\**p*<0.001, \*\*\*\**p*<0.0001.

Previous work showed that Foxo3, p53, and c-Jun can function as upstream transcriptional activators for *PUMA*, thereby promoting cell death under different cellular contexts (Fitzwalter et al., 2018; Simon et al., 2016). To test if autophagy regulates these upstream transcription activators to promote pre-OL apoptosis, we first analyzed a mutant mouse strain where *Foxo3* was conditionally deleted from oligodendrocyte lineage cells (*CNP-Cre; Foxo3^F/F^*, or *Foxo3* cKO). We found that *Foxo3* cKO mice exhibited normal myelination in the cerebellum (Figures S5E-E’), and that genetic deletion of *p53* in oligodendrocytes does not affect their survival during development (Sun *et al*., 2018). Moreover, *ATG5* cKO pre-OLs exhibited unchanged protein levels of c-Jun and phosphorylated c-Jun (Ser73) (Figure S5D), together suggesting that autophagy regulates *PUMA* mRNA levels through a novel, uncharacterized mechanism.

TFEB has been shown to be a master regulator for autophagy induction in a variety of cells upon starvation and stress (Zhao et al., 2021). A previous study revealed that TFEB promotes *PUMA/Bbc3* transcription and subsequently triggers Bax/Bak-dependent apoptosis in pre-OLs (Sun *et al*., 2018). To determine if TFEB induces autophagy in pre-OLs, we performed western blot analysis on *TFEB* control (*TFEB^F/F^*) and oligodendrocyte-specific *TFEB* knockout pre-OLs (*TFEB* KO, *Olig2-Cre; TFEB^F/F^*) with a battery of autophagy markers (Figure S5C). As expected, in the presence of BafA1 p62 and LC3-II levels were significantly increased in control pre-OLs (Figures 5G; quantified in Figure 5I). However, *TFEB* KO pre-OLs exhibited the same levels of p62 and LC3-II as compared to control cells in the presence and absence of BafA1 (Figures 5G and S5F; quantified in Figure 5I), suggesting that autophagy flux was minimally affected in *TFEB* KO pre-OLs. These observations were in strong contrast to *ATG5* cKO pre-OLs, as they showed diminished LC3-II in the presence or absence of BafA1 and that their p62 levels lost response to BafA1 treatment (Figure 5H; quantified in in Figure 5I). Finally, we found that in *ATG5* cKO pre-OLs *TFEB* mRNA and protein levels were not reduced (Figures 5D and S5B; quantified in Figure 5F), together showing that the downregulation of *PUMA* mRNA in *ATG5* cKO pre-OLs was unlikely through a TFEB-dependent manner.

To determine if autophagy functions in the TFEB-PUMA-Bax/Bak axis or acts in a separate pathway, we conducted *in vivo* genetic interaction analysis and characterized *MBP^+^* cells in the cerebellar ML among littermates with four genotypes, including the *CNP-Cre; ATG7^F/+^* (*ATG7* Het), *CNP-Cre; ATG7^F/F^* (*ATG7* KO), *CNP-Cre; TFEB^F/F^* (*TFEB* KO), and *CNP-Cre; ATG7^F/F^; TFEB^F/F^* (*ATG7;TFEB* dKO). As expected, *ATG7* KO mice exhibited significantly increased *MBP^+^* cells in the cerebellar ML as compared to *ATG7* Het mice (Left two panels in Figure 5J; quantified in Figure 5K; 1.5±0.4 cells/mm^2^ for *ATG7* Het and 11.3±2.6 cells/mm^2^ for *ATG7* KO, mean±SEM, n≥6 animals per genotype, \**p*=0.0192). *TFEB* KO mice harbored significantly increased *MBP^+^* cells in the cerebellar ML as compared to *ATG7* KO, however they were indistinguishable from *ATG7;TFEB* dKO mice (right two panels in Figure 5J; quantified in Figure 5K; 29.8±2.8 cells/mm^2^ for *TFEB* KO and 30.4±4.2 cells/mm^2^ for *TFEB;ATG7* dKO, mean±SEM, n≥3 animals per genotype, *p*=0.9981 between *TFEB* KO and *ATG7; TFEB* dKO, one-way ANOVA followed with Tukey’s multiple comparisons test). Similarly, in the cortical layer I where *ATG5* and *ATG7* cKO mice exhibited significantly increased OL numbers (Figure 2), *TFEB* cKO mice displayed enhanced phenotypes as compared to both *ATG5* and *ATG7* cKO mice, however they were indistinguishable from *PUMA^−/−^* mutants (Figure S5G). Therefore, autophagy functions in the TFEB-PUMA-Bax/Bak axis to promote pre-OL apoptosis, and it is through a noncanonical, TFEB-independent induction.

### Autophagy limits myelin wrap numbers independent of its pro-apoptotic role in pre-OLs and finetunes nerve pulse propagation

Autophagy flux remained elevated in myelinating OLs *in vivo* and *in vitro* (Figures 1 and 3). We then asked if autophagy has a separate role in those surviving pre-OLs and subsequently differentiated myelinating OLs. We analyzed the ultrastructure of myelin sheaths in the corpus callosum (CC), a white matter brain region where a great number of axons are myelinated. At P21 when CC myelination is at its peak, *ATG5* cKO mice displayed significantly increased myelin sheath thickness as compared to littermate controls, measured by *g*-ratios (the ratio of inner axonal diameter to the total outer diameter; Figures 6A-C and Figure 6G). We observed identical phenotypes in *ATG7* cKO mice (Figures 6D-F; quantified in Figures 6H and 6I) and found that these phenotypes were due to increased myelin wrap numbers (Figures 6J-K; Figures S6A and S6A’). Importantly, there is no change of myelinated axon diameter distribution in *ATG5* and *ATG7* cKO mice (Figures S6B-C’). To determine if increased myelin sheath numbers were caused by the lack of pre-OL apoptosis (Figure 2), we analyzed *CNP-Cre; Bax^F/F^;Bak^−/−^* and *PUMA^−/−^* mutants, where pre-OL apoptosis is genetically blocked (Sun *et al*., 2018). Indeed, P56 *PUMA^−/−^* mutants exhibited significantly increased OL numbers in the CC (Figure S6I). However, neither *CNP-Cre; Bax^F/F^;Bak^−/−^* nor *PUMA^−/−^* mutants displayed alterations in myelin sheath thickness (Figures 6L-N; Figures S6F-H), myelin wrap number distribution (Figures S6D-D’), or myelinated axon diameter distribution (Figures S6E-E’). These results showed that aberrantly increasing OL number by blocking pre-OL apoptosis does not contribute to myelin sheath thickness. Thus, autophagy continuously functions in surviving pre-OLs and myelinating OLs to limit myelin wrap numbers and sheath thickness independent of its pro-apoptotic role in pre-OLs.

**Figure 6.**
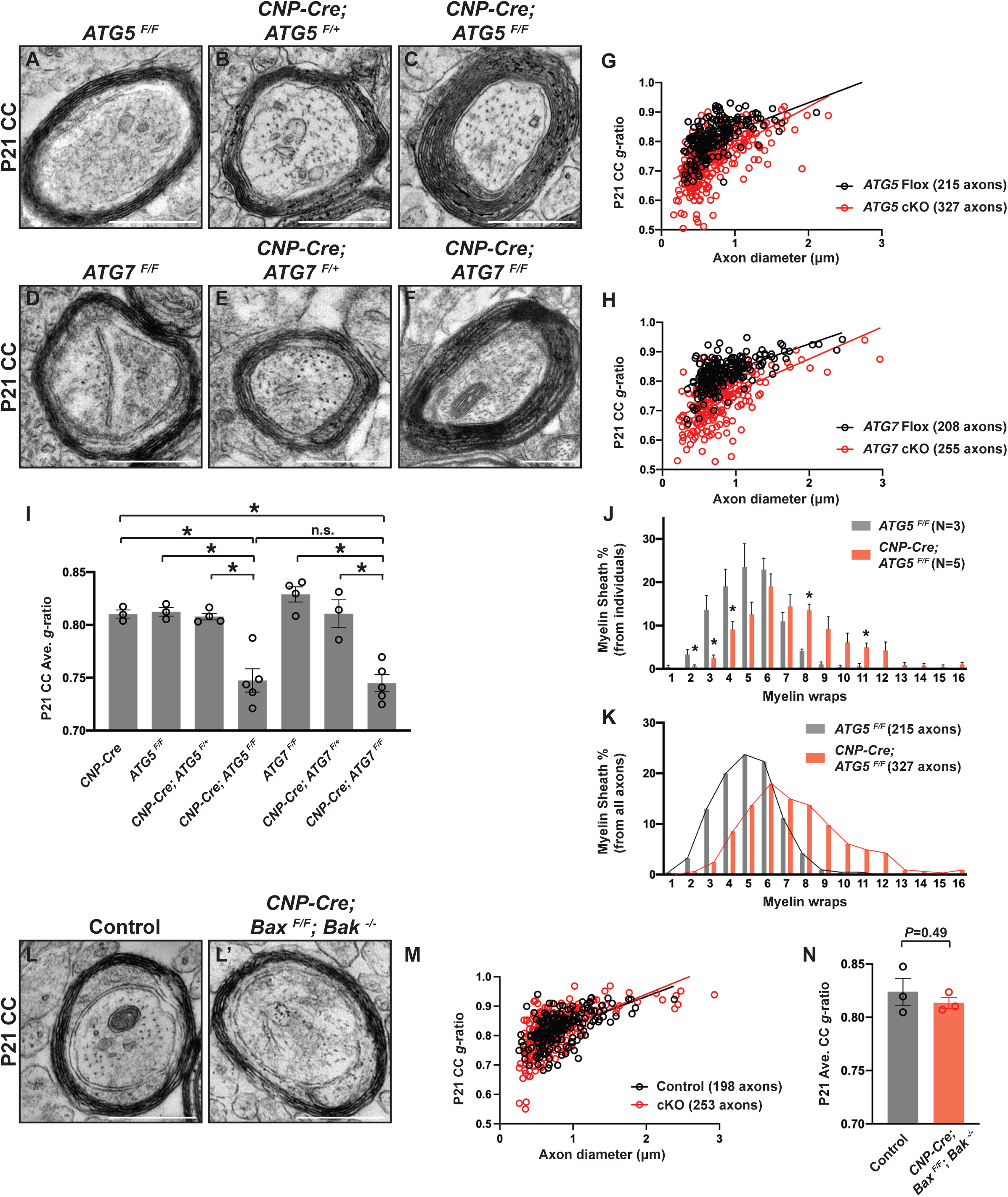
Autophagy restricts myelin wrap numbers and sheath thickness independent of its pro-apoptotic roles in the pre-OLs. **(A-F)** Representative TEM micrographs from P21 *ATG5^F/F^* (A), *CNP-Cre; ATG5^F/+^* (B), *CNP-Cre; ATG5^F/F^* (C), *ATG7^F/F^* (D), *CNP-Cre; ATG7^F/+^* (E), and *CNP-Cre; ATG7^F/F^* (F) corpus callosum (CC), showing that *CNP-Cre; ATG5^F/F^* (C) and *CNP-Cre; ATG7^F/F^* mutants (F) exhibited increased myelin sheath thickness. **(G and H)** Scatter plot showing *g*-ratios of myelinated axons as a function of axon diameter in P21 *CNP-Cre; ATG5^F/F^* (*ATG5* cKO) and *ATG5^F/F^* (*ATG5* Flox) mice (G), and the scatter plot showing *g*-ratios between P21 *CNP-Cre; ATG7^F/F^* (*ATG7* cKO) and *ATG7 ^F/F^* (*ATG7* Flox) mice (H). Both *ATG5* and *ATG7* cKO mice exhibited reduced *g*-ratios among axons with diameters <1 μm. **(I)** Quantification of average *g*-ratios of myelinated axons in P21 *CNP-Cre*, *ATG5^F/F^*, *CNP-Cre; ATG5^F/+^*, *CNP-Cre; ATG5^F/F^*, *ATG7^F/F^*, *CNP-Cre; ATG7^F/+^*, and *CNP-Cre; ATG7^F/F^* corpus callosum, showing that the *CNP-Cre; ATG5^F/F^* and *CNP-Cre; ATG7^F/F^* mutants exhibited significant reduced *g*-ratios. **(J and K)** Quantifications of averaged percentage of myelinated axons with different myelin wraps from individual animals (J) or total percentage of myelinated axons with different myelin wraps (K) between P21 *ATG5^F/F^* (Flox) and *CNP-Cre; ATG5^F/F^* (cKO) mice. *CNP-Cre; ATG5^F/F^* mutants exhibited a right-shift distribution of myelin wrap percentage, showing that the decreased *g*-ratios observed in the *CNP-Cre; ATG5^F/F^* mutants (G) were due to increased myelin wrap numbers. **(L and L’)** Representative TEM micrographs of P21 corpus callosum from *CNP-Cre; Bax^F/F^; Bak^−/−^* mutants (L’) and littermate controls (L, *Bax^F/F^;Bak^−/−^*). **(M and N)** Quantifications of average *g*-ratios of myelinated axons (N), and as a function of axon diameter (M), in P21 *CNP-Cre; Bax^F/F^; Bak^−/−^* mutants (red) and littermate controls (black).

To determine how the hypermyelination phenotype observed in the *CNP-Cre; ATG5^F/F^* mutants impacts neuronal activity and nerve impulse propagation, we measured the compound action potentials (CAPs) on the corpus callosum (Figure 7A) (Li et al., 2016; Reeves et al., 2005). In response to stimulation, both control (*CNP-Cre; ATG5^F/+^*) and *ATG5* cKO mice (*CNP-Cre; ATG5^F/F^*) exhibited two distinct peaks, N1 and N2 peaks, representing fast and slow action potential propagation, respectively (Figure 7B). Surprisingly, we found that the latencies of N1 and N2 remained unchanged in *ATG5* cKO mice at various recording locations except that the N2 latency at 0.5 mm showed a longer latency in *ATG5* cKO mice (Figures 7C and 7D, left panels). Consistent with this observation, the conduction velocity showed no significant difference between groups (Figures 7C and 7D, right panels). However, we found that the sizes of N2 peaks were significantly reduced in *ATG5* cKO mice (1V stimulation, N2=1.140 ± 0.120 mV•ms for control, and N2=0.808 ± 0.095 mV•ms for *ATG5* cKO, n = 30 brain slices from 10 control mice and n=26 slices from 10 *ATG5* cKO mice; see quantification in the left panel of Figure 7F), whereas N1 peaks were unchanged (1V stimulation, N1=0.962 ± 0.055 mV•ms for control, and N1=1.021 ± 0.051 mV•ms for *ATG5* cKO; see quantification in the left panel of Figure 7E). These deficits were further revealed by the disrupted N1/N2 balance in *ATG5* cKO mice, as they exhibited a reduced N2 ratio and an increased N1 ratio (right panels in Figures 7E and 7F; N2 ratio = Area_N2_/(Area_N1+N2_), 0.518 ± 0.020 for control and 0.417 ± 0.026 for *ATG5* cKO; N1 ratio = Area_N1_/(Area_N1+N2_), 0.483 ± 0.020 for control and 0.583 ± 0.026 for *ATG5* cKO). This data showed that although the conduction velocity of CAP was not affected, *ATG5* cKO mice exhibited an increase in the sum of fast propagating CAP and a decrease in the sum of slow propagating CAP. It also indicates that the balance between fast- and slow-propagation of impulses along the corpus callosum was altered in *ATG5* cKO mice. Taken together, autophagy promotes apoptosis in subsets of premyelinating oligodendrocytes to control myelination spatiotemporal specificity, and it continuously functions in myelinating oligodendrocytes to limit myelin wrap numbers and regulates critical aspects of action potential propagation.

**Figure 7.**
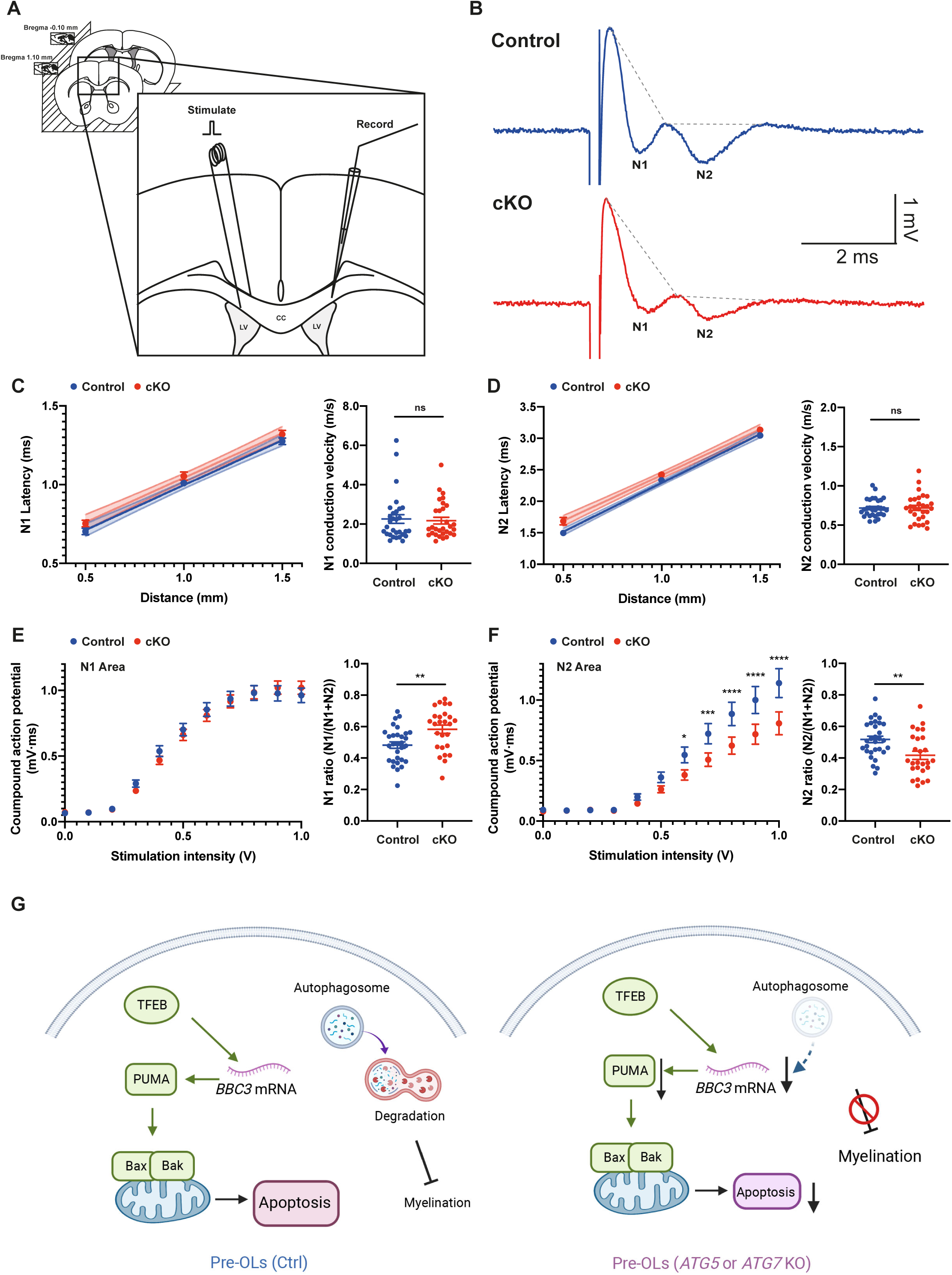
*CNP-Cre; ATG5^F/F^* mutants exhibit aberrant compound action potentials. **(A)** Schematics of compound action potentials (CAPs) recording. To measure the conduction along the corpus callosum, the CAPs were measured at various distance from 0.5 mm to 1.5 mm between stimulus and recording sites. **(B)** Representative traces of CAPs recorded on the corpus callosum from control (*CNP-Cre; ATG5^F/+^*) and *ATG5* cKO mice (*CNP-Cre; ATG5^F/F^*) in response to 1 V stimulation at 1 mm distance. The dashed line represents the margin of peak areas. **(C and D)** The latency of N1 and N2 peaks were measured at difference distance between stimuli and recording sites and linear regression analysis was performed. In linear regression analysis, N1 and N2 latency showed a similar pattern without the significant difference in the slop (left panels in C and D, respectively; N1, *p* = 0.9203; N2, *p* = 0.2018). The conduction velocity of N1 (right panel in C) and N2 (right panel in D) were calculated with the equation: (latency_1.5mm_ - latency_1mm_)/0.5mm. **(E and F)** Plotting area of CAPs responding to various stimulation intensity (from 0 V to 1 V) for N1 peak (E) and N2 peak (F). N1 ratio and N2 ratio were calculated by Area_N1_/(Area_N1+N2_) and Area_N2_/(Area_N1+N2_) at 1 V stimuli, respectively. Two-way ANOVA and *t*-test was used to test statistical significance. **(G)** Autophagy functions cell-autonomously to promote apoptosis in subsets of premyelinating oligodendrocytes, and it continuously functions in myelinating oligodendrocytes to limit myelin wrap numbers and to control critical aspects of nerve pulse propagation. In control animals, the TFEB-PUMA-Bax/Bak pathway drives apoptosis in subsets of pre-OLs, resulting in elimination of a great faction of pre-OLs during development. In the absence of ATG5 or ATG7, autophagy flux is perturbed, which further leads to reduced *PUMA/Bbc3* mRNA levels. Autophagy-dependent PUMA protein reduction decreases pre-OL apoptosis and disrupts myelination spatiotemporal specificity. Autophagy continuously functions in the surviving pre-OLs and myelinating OLs to limit myelin wrap numbers and myelin sheath thickness independent of its pro-apoptotic roles in pre-OLs, thereby regulating critical aspects of nerve pulse propagation. Error bars represent SEM. The significance was presented as asterisk. \**p*< 0.05, \*\**p*< 0.01, \*\*\**p*< 0.001, \*\*\*\**p*<0.0001.

## DISCUSSION

The fine balance between construction and destruction of cells governs organogenesis, tissue remodeling, cancer, and aging (Green, 2019). Autophagy and apoptosis pathways intimately interact with each other to determine cell fate. Despite the evidence showing that autophagy can promote apoptosis in invertebrates, it remains controversial if it plays similar functions in vertebrates under physiological conditions. In this study, we revealed that autophagy promotes apoptosis in subsets of immature myelinating glia (premyelinating oligodendrocytes) and regulates nerve impulse propagation, thereby ensuring spatiotemporal specificity and functional integrity of CNS myelination. We found that autophagy flux is elevated at the premyelinating OL stage during differentiation, and that it functions cell-autonomously to induce apoptosis in subsets of pre-OLs. Genetic blockage of autophagy flux by conditionally deleting *ATG5* or *ATG7* in oligodendrocyte lineage cells leads to ectopic pre-OL survival, which further causes aberrant myelination in the unmyelinated brain regions, increased oligodendrocyte numbers across diverse brain areas, as well as altered myelination timing. We found that autophagy genetically interacts with the TFEB-PUMA-Bax/Bak apoptotic pathway and that autophagy elevates *PUMA* mRNA levels to trigger pre-OL apoptosis. Finally, autophagy continuously functions in the myelinating OLs to limit myelin sheath thickness independent of its pro-apoptotic role in pre-OLs, thereby patterning CNS myelination during development (Figure 7G).

Under nutrient deprivation and stress conditions, induction of autophagy promotes cell adaption and survival. Autophagy-associated cell death remains rare and is mostly observed in invertebrates. Our study showed that in the absence of *ATG5* or *ATG7*, two critical genes mediating autophagosome elongation and autophagy flux, subsets of pre-OLs no long undergo apoptosis *in vitro* (Figure 4) and *in vivo* (Figure 2). Consequently, myelination location and timing are severely disrupted when those pre-OLs ectopically survive. These observations strongly showed that autophagy promotes cell death in a specialized glial cell type under physiological conditions and that autophagy-mediated pre-OL cell death is critical for normal nervous system development. Importantly, our data revealed that autophagy-mediated pre-OL cell death is indeed apoptosis rather than other forms of cell death, including autosis and necrosis (Figure 4). Together these results provide key evidence showing how autophagy collaborates with apoptosis pathways to promote programmed cell death in mammalian cells under physiological relevant circumstances.

What are the molecular mechanisms underlying autophagy-mediated pre-OL apoptosis? Our data showed that autophagy genetically interacts with the TFEB-PUMA-Bax/Bak pathway (Figure 5), a pathway that powerfully promotes pre-OL apoptosis (Sun *et al*., 2018). Intriguingly, both *in vitro* and *in vivo* phenotypes of autophagy-deficient pre-OLs are milder than the ones shown by *TFEB* cKO and *PUMA^−/−^* mutants (Figure S5) (Sun *et al*., 2018). In addition, there is no enhancement of the phenotype in *ATG7; TFEB* double knockout mice as compared to *TFEB* cKO mice (Figure 5), suggesting that the TFEB-PUMA-Bax/Bak apoptotic pathway is the main driver for pre-OL cell death while autophagy plays a modulatory role (Figure 7G). Moreover, autophagy is not induced by TFEB under the basal condition during oligodendrocyte differentiation, as there are minimal autophagy flux changes in *TFEB* cKO pre-OLs (Figure 5). This is not surprising since TFE3, another master regulator for autophagy induction, is also expressed by pre-OLs and thus it could play a redundant role in the absence of TFEB (Sun *et al*., 2018; Zhao *et al*., 2021). In addition, recent work showed that in zebrafish microglia *Tfeb* and *Tfe3* are dispensable for basal levels of autophagy and lysosomal gene expression under homeostatic conditions (Iyer et al., 2022). Consistent with these observations, we found that autophagy elevates *PUMA* mRNA levels through a TFEB-independent manner (Figure 5). Together these results suggest that autophagy functions upstream of *PUMA*, a strong inducer for apoptosis, however it acts independent of TFEB-mediated induction (Figure 7G).

Previous work showed that p53, Foxo3, and c-Jun are transcriptional activators for *PUMA/Bbc3* (Fitzwalter *et al*., 2018; Simon *et al*., 2016). However, oligodendrocyte-specific deletion of p53 or Foxo3 does not cause ectopic pre-OL survival (Figure S5) (Sun *et al*., 2018). Additionally, c-Jun protein levels as well as phosphorylated c-Jun (Ser73) remain unchanged in the absence of ATG5 (Figure S6), together showing that autophagy regulates *PUMA* mRNA through a novel, uncharacterized mechanism. Because *PUMA* mRNA levels are decreased in the absence of ATG5 and because autophagy’s major function is to degrade cytoplasm contents, autophagy might degrade transcriptional inhibitors for *PUMA* to enhance its mRNA expression. Alternatively, ERphagy and ER stress pathways may act upstream of *PUMA* to control pre-OL fate, given that our quantitative mass spectrometry analysis revealed that ERphagy receptors are aberrantly upregulated in *ATG5* cKO pre-OLs (Figure 5) and that the activation of ER stress pathways can induce OL apoptosis (Lin and Popko, 2009). It will be of great interest to determine the molecular mechanisms linking autophagy with *PUMA* mRNA expression.

After pre-OLs differentiate into mature OLs, they drastically expand their plasma membrane over 6,000 folds within a few days, making them the most powerful cell type in the CNS to generate lipid and lipid-associated proteins (Snaidero and Simons, 2014). Our results showed that autophagy flux remains elevated in mature OLs (Figures 1 and 3), and that autophagy continuously functions in the surviving pre-OLs and myelinating OLs to limit myelin wrap numbers during development (Figure 6). A previous study found that conditionally deleting *ATG5* in OPCs via an inducible manner causes OL cell death, hypomyelination, and lethality early postnatally (Bankston et al., 2019). However, our work demonstrated that four conditional knockout mouse strains with genetic perturbation of autophagy in oligodendrocyte lineage cells are viable and fertile, including the *CNP-Cre; ATG7^F/F^*, *CNP-Cre; ATG5^F/F^*, *Olig2-Cre; ATG5^F/F^*, and *PDGFRα-Cre^ERT2^; ATG5^F/F^* mutant mice. In fact, we found opposite phenotypes in these mutants where all of them exhibit increased OL numbers across diverse brain regions. Morever, both *CNP-Cre; ATG5^F/F^* and *CNP-Cre; ATG7^F/F^* mutants show identical hypermyelination phenotypes in the corpus callosum (Figure 6). Our results are consistent with the recent finding showing that *ATG7* cKO mice are viable and hypermyelinated (Aber et al., 2022). Therefore, future work is needed to delineate differentiation-stage-specific function of autophagy and the underlying mechanisms to fully understand its roles in the establishment and maintenance of myelination.

It is well established that hypomyelination severely disrupts nervous system function by greatly slowing down nerve impulse propagation and depriving trophic support for axons (Lehman and Harrison, 2002; Moore et al., 2020). Conversely, it remains poorly understood how hypermyelination, either by harboring excessive oligodendrocytes and/or by aberrantly thickening myelin sheaths, affects nervous system function (Yu et al., 2011). We found that the conduction velocity of CAP in the CC is not changed in *ATG5* cKO mice, but the fine balance between slow-versus fast-conducting nerves is altered. It remains unclear how hypermyelination leads to this physiological defect, and how this phenotype impacts callosal axons with different calibers. Together with a recent work that linked aberrant neural activity (seizures) with hypermyelination and hyper-synchronization in thalamocortical network (Knowles et al., 2022), it will be of great importance to determine the precise roles of oligodendrocyte number, myelin sheath thickness, as well as oligodendrocyte metabolism in modulating nerve pulse propagation in a circuit-specific manner.

In conclusion, autophagy promotes premyelinating oligodendrocyte apoptosis to control the location and timing of CNS myelination during development. Our observations provide a mechanistic link connecting autophagy and apoptosis, two evolutionarily conserved cellular processes that intimately interact with each other to determine cell fate. It will be of great interest to address if the fine balance between these two pathways within myelinating glia regulates myelin degeneration and regeneration.

## ACKNOWLEGEMENTS

We thank Dr. Klaus-Armin Nave in sharing the *CNP-Cre* mice. We thank Drs. John Abrams, Yang Liu, and Helmut Kramer for helpful comments on the manuscript. We also thank Dr. David J. Simon and members of Sun laboratory for assistance and discussions. This work was supported by UT Southwestern Endowed Scholarship (L.O.S.), NIH R00EY029330 (L.O.S.), Texas Alzheimer’s Research and Care Consortium (L.O.S.), Brain & Behavior Research Foundation (L.O.S.), Welch Foundation (L.O.S.), DoD W81XWH-21-1-0830 (L.O.S.), NIH 1DP2MH129988 (L.O.S.), 1S10OD021685-01A1 (UT Southwestern electron microscopy core facility), and NIH R01DC03157 (K.J.H.). L.O.S. is a Southwestern Medical Foundation Scholar in Biomedical Research and the John and Polly Sparks Foundation Investigator.

## AUTHOR CONTRIBUTIONS

Conceptualization, K.J.H., and L.O.S.; Methodology, T.Z., H.G.B., A.B., Y.Z., D.B., K.J.H., and L.O.S.; Investigation, T.Z., H.G.B., A.B., Y.Z., D.B., J.X., S.W. S.B.M., K.J.H., and L.O.S.; Writing, K.J.H. and L.O.S.; Funding Acquisition, K.J.H. and L.O.S.; Supervision, K.J.H. and L.O.S..

## DECLRATION OF INTERETS

None.

## SUPPLEMENTAL INFORMATION

### Supplemental Figures

**Figure S1, related to Figure 1. Characterization of pre-OLs and mature OLs *in vitro* by confocal and transmission electron microscopy.**

**Figure S2, related to Figure 2. Deletion of *ATG5* or *ATG7* in oligodendrocyte lineage cells disrupts the location and timing of CNS myelination.**

**Figure S3, related to Figure 3. Characterization of autophagosomes in differentiating oligodendrocytes by transmission electron microscopy (TEM).**

**Figure S4, related to Figure 4. Characterization of OPC differentiation and pre-OL cell death in the absence of ATG5.**

**Figure S5, related to Figure 5. Biochemical characterization of *TFEB*- and *ATG5*-deficient oligodendrocytes and analysis of excessive oligodendrocytes in autophagy and apoptosis mutant mice.**

**Figure S6, related to Figure 6. Characterization of axon diameter and myelin sheath number in the *CNP-Cre; ATG7^F/F^*, *CNP-Cre; ATG5^F/F^, CNP-Cre; Bax^F/F^; Bak^−/−^*, and *PUMA^−/−^* mutant mice.**

### Other Supplemental Items

**Supplemental table 1 (Table S1): Bulk RNA sequencing analysis of *in vivo* autophagy gene expression (GO:0006914) in seven major brain cell types, related to Figure 1.** *In vivo* Autophagy gene mRNA expression levels (average FKPM) were shown in astrocytes (Astro), neurons (N), oligodendrocyte precursor cells (OPC), premyelinating oligodendrocytes (pre-OL), myelinating oligodendrocytes (OL), microglia (Micro), and endothelia cells (Endo) from a published dataset (Zhang *et al*., 2014).

**Supplemental table 2 (Table S2): Quantitative mass spectrometry analysis of dysregulated proteins in *CNP-Cre; ATG5^F/F^* (*ATG5* cKO) pre-OLs as compared to *CNP-Cre; ATG5^F/+^* (*ATG5* Het) pre-OLs, related to Figure 5.** Two tabs were shown including (from left to right): All 5,557 detected proteins in *ATG5* Het and cKO pre-OLs ranked by log2 fold change, and proteins that were significantly up- or down-regulated (log2 fold change >0.6 or <-.06). The criterion for statistical significance was set at *p*<0.05.

**Supplemental table 3 (Table S3): Differential gene expression analysis between *ATG5* cKO pre-OLs and *ATG5* Het pre-OLs, Related to Figure 5.** One tab showing gene expression measured by normalized Htseq-counts (ranked by *p* values) and differential gene expression analysis between *ATG5* cKO pre-OLs and *ATG5* Het pre-OLs by DEseq2. The criterion for statistical significance was set at *p*<0.05.

**Supplemental movie 1 (Movie S1): Live-cell imaging of differentiating oligodendrocytes using OPCs purified from P10 *CNP-Cre; ATG5^F/+^* mice, related to Figure 4.** IncuCyte Annexin V red reagent was applied in the culture medium at the beginning of imaging and supplemented every two days throughout the entire course of imaging (5 days of total live-cell imaging with one image being taken every 2 hours). The video is displayed at 4 frames per second.

**Supplemental movie 2 (Movie S2): Live-cell imaging of differentiating oligodendrocytes using OPCs purified from P10 *CNP-Cre; ATG5^F/F^* mice, related to Figure 4.** IncuCyte Annexin V red reagent was applied in the culture medium at the beginning of imaging and supplemented every two days throughout the entire course of imaging (5 days of total live-cell imaging with one image being taken every 2 hours). The video is displayed at 4 frames per second. In comparison to *CNP-Cre; ATG5^F/+^* oligodendrocytes, *CNP-Cre; ATG5^F/F^* mutant oligodendrocytes exhibited reduced cell death during differentiation. See quantification in Figure 4.

## SUPPLEMENTAL FIGURE LEGENDS

**Figure S1, related to Figure 1.**
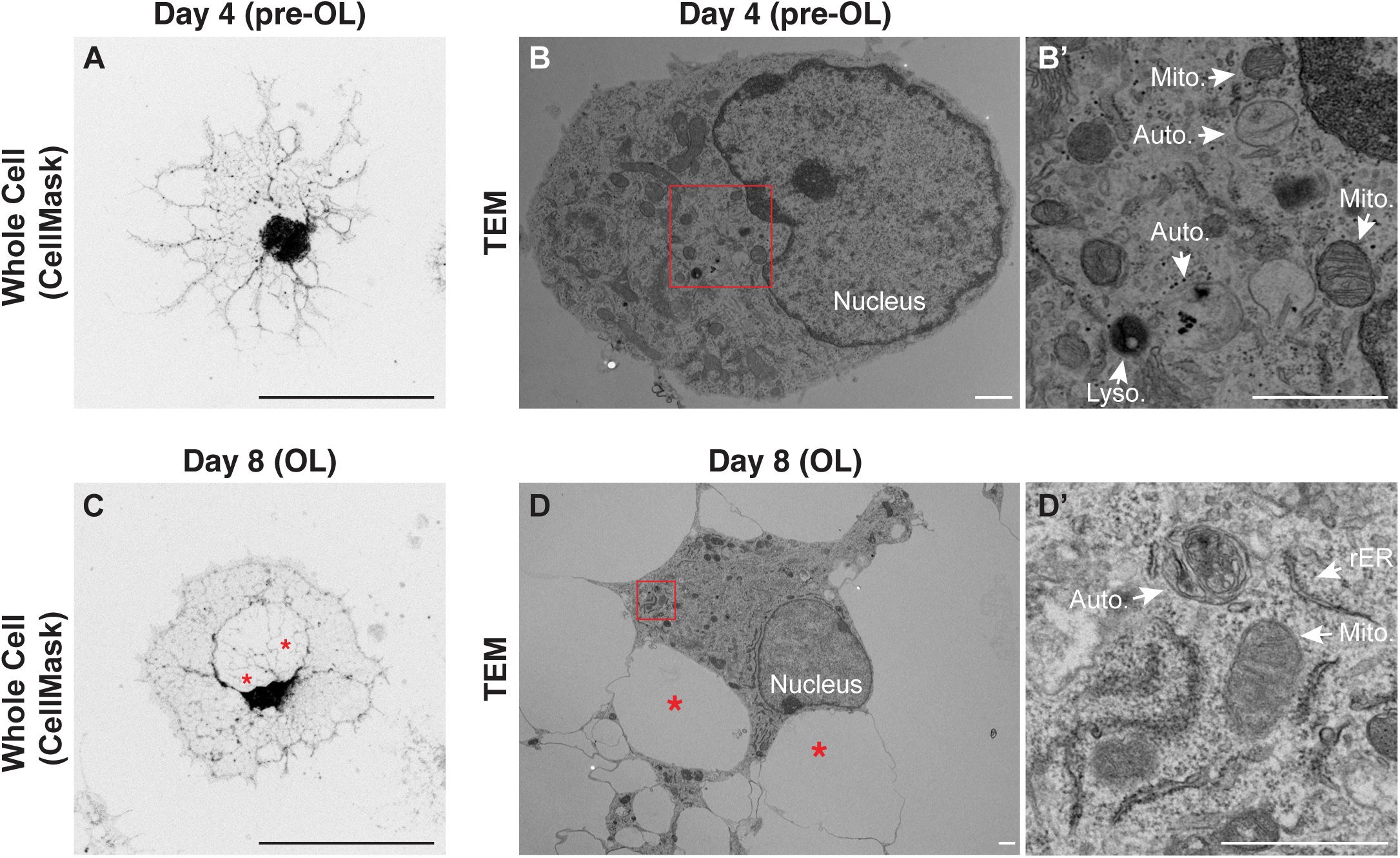
Characterization of pre-OLs and mature OLs *in vitro* by confocal and transmission electron microscopy. **(A)** Representative confocal micrograph of a premyelinating oligodendrocyte (pre-OL) at differentiation day 4 counterstained by CellMask. **(B and B’)** Representative transmission electron microscopy (TEM) micrograph of a pre-OL at differentiation day 4. B’ represents the inset in B. **(C-D’)** Representative confocal (C) and TEM micrographs (D) of mature oligodendrocytes (OLs) at differentiation day 8. D’ represents the inset in D. Mito., mitochondria. Auto., autophagosome. Lyso., lysosome. rER, rough endoplasmic reticulum. Asterisks indicate the gaps formed between major branches of mature OLs by confocal microscopy (C) and TEM (D). See more examples and quantification in Figures 3 and S3. Scale bars: 100 μm in (A) and (C); 1 μm in (B), (B’), (D), and (D’).

**Figure S2, related to Figure 2.**
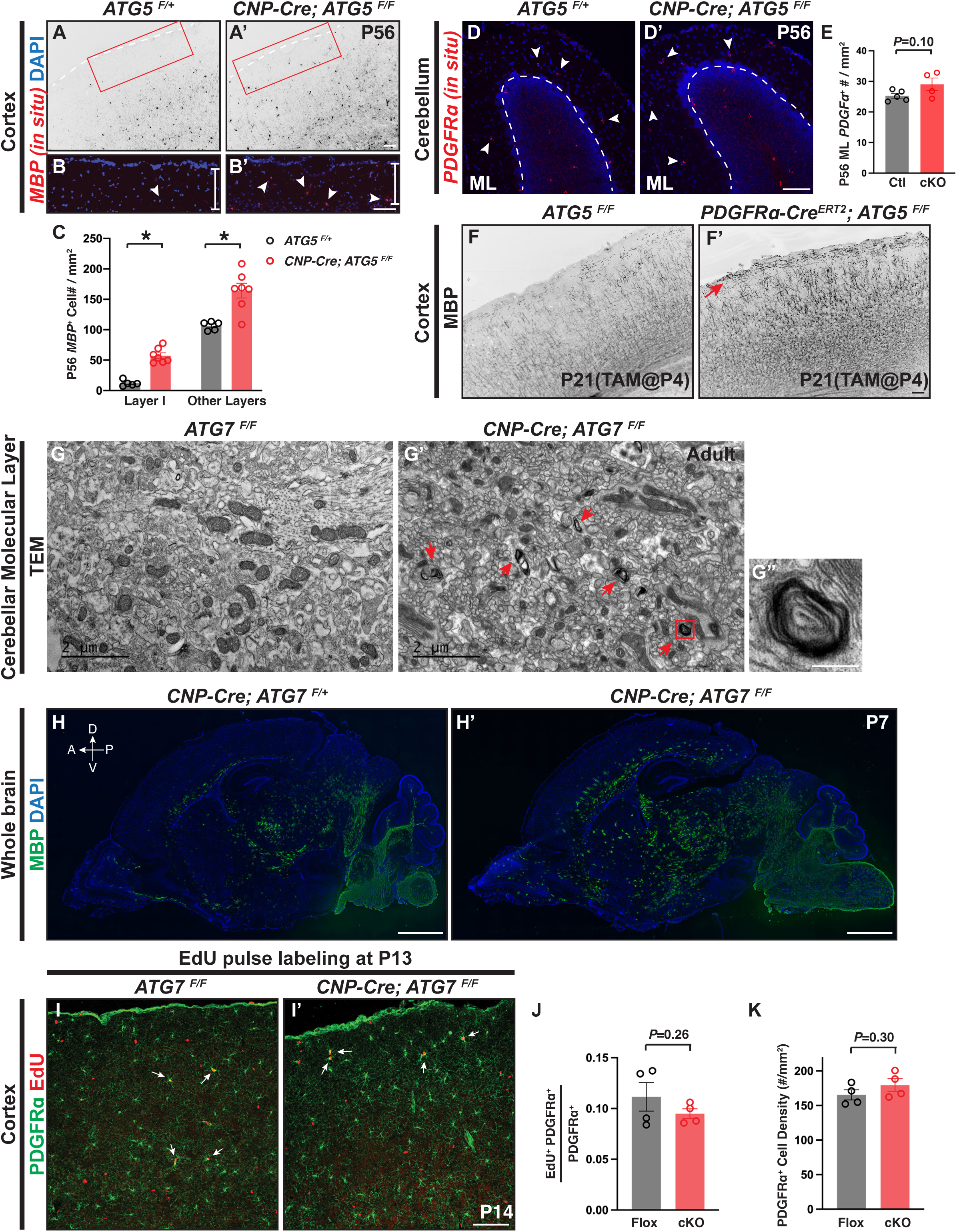
Deletion of *ATG5* or *ATG7* in oligodendrocyte lineage cells disrupts the location and timing of CNS myelination. **(A-B’)** Representative micrographs of P56 *ATG5^F/+^* (A and B) and *CNP-Cre; ATG5^F/F^* (A’ and B’) sagittal brain sections labeled by *MBP in situ* hybridization probes. White arrows in B and B’ indicate *MBP^+^* oligodendrocytes in cortical layer I. B and B’ represent the insets in A and A’, respectively. White dashed lines in A and A’ demarcate the cortical pia surface. **(C)** Quantification of *MBP^+^* oligodendrocytes in cortical layer I and other cortical layers in P56 *ATG5^F/+^* and *CNP-Cre; ATG5^F/F^* mice, showing that *CNP-Cre; ATG5^F/F^* mutants exhibited significantly increased *MBP^+^* cell numbers in the cortex. **(D and D’)** Representative confocal micrographs of P56 *ATG5^F/+^* and *CNP-Cre; ATG5^F/F^* cerebellum labeled by *PDGFRα in situ* probes (red). White arrowheads indicate *PDGFRα*^+^ OPCs in the cerebellar molecular layer (ML). **(E)** Quantification of *PDGFRα*^+^ cell density in the cerebellar ML of P56 control (ctl, *ATG5^F/+^*) and cKO (*CNP-Cre; ATG5^F/F^*). **(F and F’)** Representative micrographs of *ATG5^F/F^* (F) and *PDGFRα-Cre^ERT2^; ATG5^F/F^* (F’) brain sections that were injected with tamoxifen at P4 and analyzed at P21. Red arrow indicates excessive MBP immunolabeling in cortical layer I. n=3 animals per genotype. **(G-G”)** Representative TEM micrographs of *ATG7^F/F^* (G) and *CNP-Cre; ATG7^F/F^* (G’) cerebellar molecular layer. Red arrows in G’ indicate aberrant myelin wrapping in *CNP-Cre; ATG7^F/F^* cerebellar ML. G” represents the inset in G’. n=2 animals per genotype. **(H and H’)** Representative tile-scanned micrographs of P7 *CNP-Cre; ATG7^F/F^* (H’) and littermate control (*CNP-Cre; ATG7^F/+^* in H) immunostained with the antibody raised against MBP (green), showing that *CNP-Cre; ATG7^F/F^* mutants exhibited precocious MBP immunoreactivity across diverse brain regions. n=3 animals per genotype. A, anterior; P, posterior; D, dorsal; V, ventral. **(I and I’)** Representative confocal micrographs of *ATG7^F/F^* (I) and *CNP-Cre; ATG7^F/F^* (I’) cortices pulse-labeled by 5-ethynyl-2’-deoxyuridine (EdU; red) at P13 and characterized 24 hours later with the antibody raised against PDGFRα (green). White arrows indicate PDGFRα^+^ and EdU^+^ double positive cells. **(J)** Quantification of OPC proliferation as indicated by the ratio of EdU^+^; PDGFRα^+^ double positive cells over total PDGFRα^+^ cells in *ATG7^F/F^* (Flox) and *CNP-Cre; ATG7^F/F^* (cKO) cortices. **(K)** Quantification of PDGFRα^+^ cell density in P14 *ATG7^F/F^* (Flox) and *CNP-Cre; ATG7^F/F^* (cKO) cortices. Error bars represent SEM. Scale bars: 100 μm in (A’) for (A) and (A’); 100 μm in (B’) for (B) and (B’); 100 μm in (D’) for (D) and (D’); 100 μm in (F’) for (F) and (F’); 2 μm in (G) and (G’); 200 nm in (G”); 1 mm in (H’) for (H) and (H’); and 100 μm in (I’) for (I) and (I’). \**p*<0.05.

**Figure S3, related to Figure 3.**
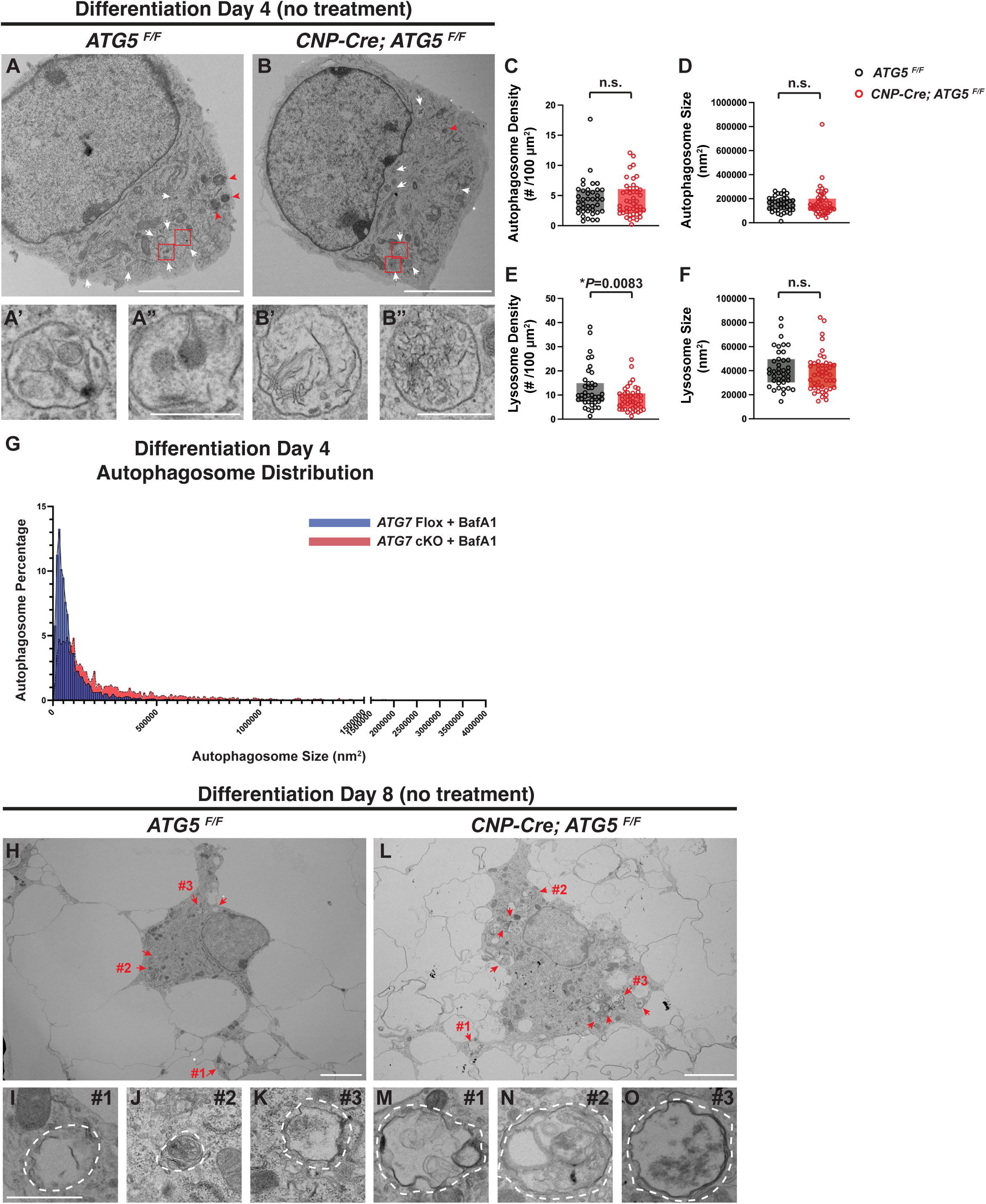
Characterization of autophagosomes in differentiating oligodendrocytes by transmission electron microscopy (TEM). **(A-B”)** Representative TEM micrographs of *ATG5^F/F^* (A-A”) and *CNP-Cre; ATG5^F/F^* (B-B”) at differentiation day 4. A’-A” and B’-B” represent enlarged views of red insets in A and B, respectively, showing autophagosomes in *ATG5^F/F^* (A’-A”) and *CNP-Cre; ATG5^F/F^* pre-OLs (B’-B”). Red arrowheads: lysosomes. White arrows: autophagosomes. **(C-F)** Quantifications (box and whisker, min to max) of autophagosome density (C), autophagosome size (D), lysosome density (E), and lysosome size (F) in *ATG5^F/F^* and *CNP-Cre; ATG5^F/F^* oligodendrocytes at differentiation day 4. n≥39 cells per genotype. **(G)** Distribution curves of autophagosomes with different sizes in *ATG7^F/F^* (*ATG7* flox, blue) and *CNP-Cre; ATG7^F/F^* (*ATG7* cKO, red) oligodendrocytes in the presence of Bafilomycin A1(BafA1) treatment. *ATG7* cKO oligodendrocytes exhibited increased percentages of large autophagosomes as compared to *ATG7* Flox oligodendrocytes, as indicated by the right shift of its distribution curve. n≥20 cells per genotype. **(H-O)** Representative TEM micrographs of *ATG5^F/F^* (H-K) and *CNP-Cre; ATG5^F/F^* (L-O) at differentiation day 8, showing that *CNP-Cre; ATG5^F/F^* mutant oligodendrocytes harbored aberrantly enlarged autophagosomes. I-K on the bottom left represent the enlarged views of autophagosomes in H as indicated by #1-3, respectively. M-O on the bottom right represent the enlarged views of autophagosomes in L as indicated by #1-3, respectively. Red arrows: autophagosomes. White dashed lines demarcate autophagosomes. Scale bars: 5 μm in (A) and (B); 1 μm in (A”) for (A’) and (A”); 1 μm in (B”) for (B’) and (B”); 5 μm in (H) and (L); and 1 μm in (I) for (I)-(O). \**p*<0.05.

**Figure S4, related to Figure 4.**
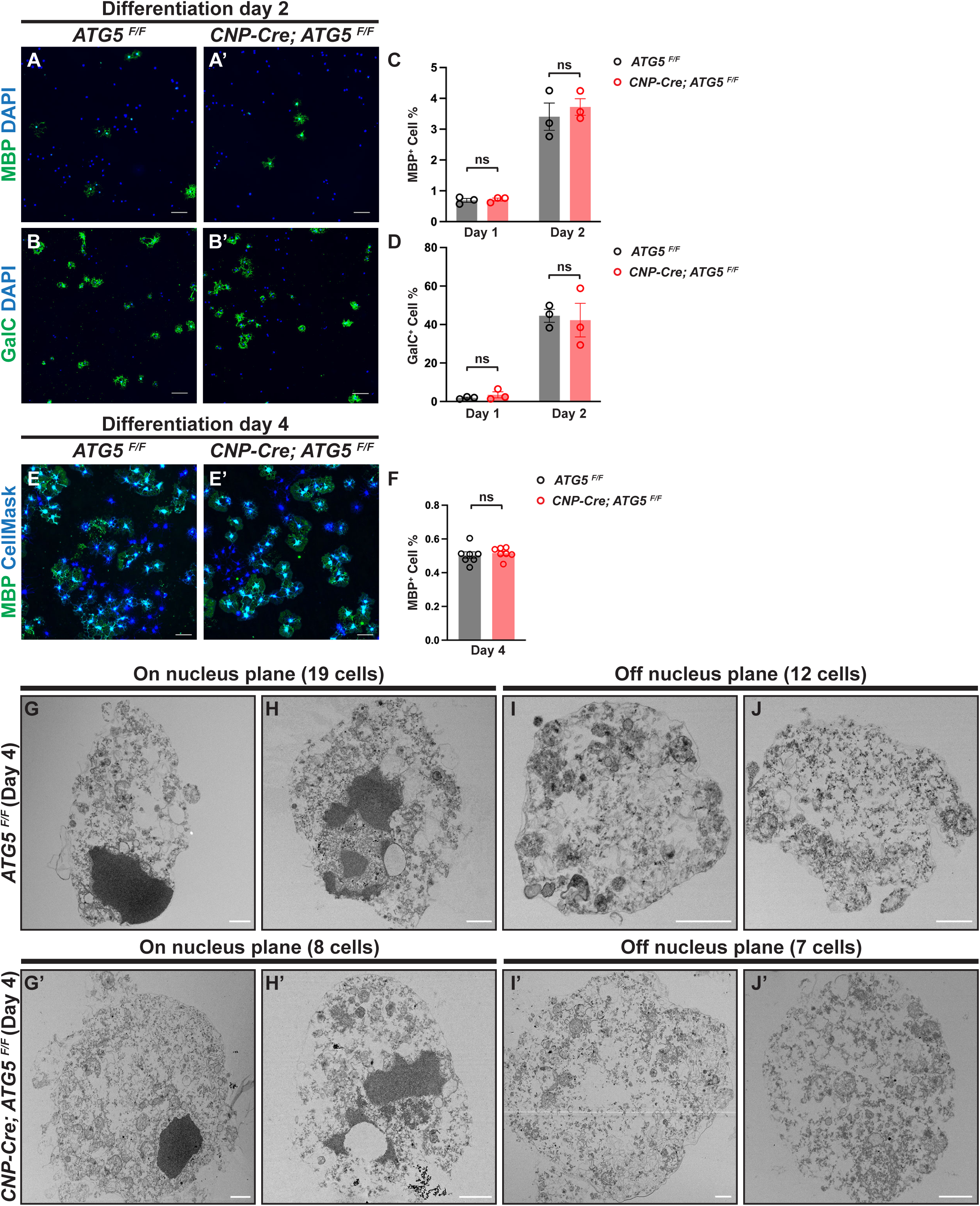
Characterization of OPC differentiation and pre-OL cell death in the absence of ATG5. **(A-D)** Representative micrographs of *ATG5^F/F^* (A and B) and *CNP-Cre; ATG5^F/F^* oligodendrocytes (A’ and B’) at differentiation day 2 labeled by antibodies raised against MBP (green in A and A’) or GalC (green in B and B’). The percentages of MBP^+^ cells and GalC^+^ cells were quantified in C and D, respectively. **(E and E’)** Representative micrographs of *ATG5^F/F^* (E) and *CNP-Cre; ATG5^F/F^* oligodendrocytes (E’) at differentiation day 4 stained by an antibody raised against MBP (green) and counterstained by CellMask (blue). **(F)** Quantification of MBP^+^ oligodendrocyte percentage at differentiation day 4 in control and *ATG5* cKO oligodendrocytes. **(G-J’)** Representative TEM micrographs of dead oligodendrocytes from *ATG5^F/F^* (G-J) and *CNP-Cre; ATG5^F/F^* mice (G’-J’) with the sections on the nucleus plane (G, H, G’, and H’) and the ones off the nucleus plane (I, J, I’, and J’), showing that a small subset of *CNP-Cre; ATG5^F/F^* still underwent apoptosis with the characteristics of nucleus condensation and intact plasma membrane. Error bars represent SEM. Scale bars: 100 μm in (A), (B), (A’), (B’), (E), and (E’); 1μm in (G)-(J’). \**p*<0.05.

**Figure S5, related to Figure 5.**
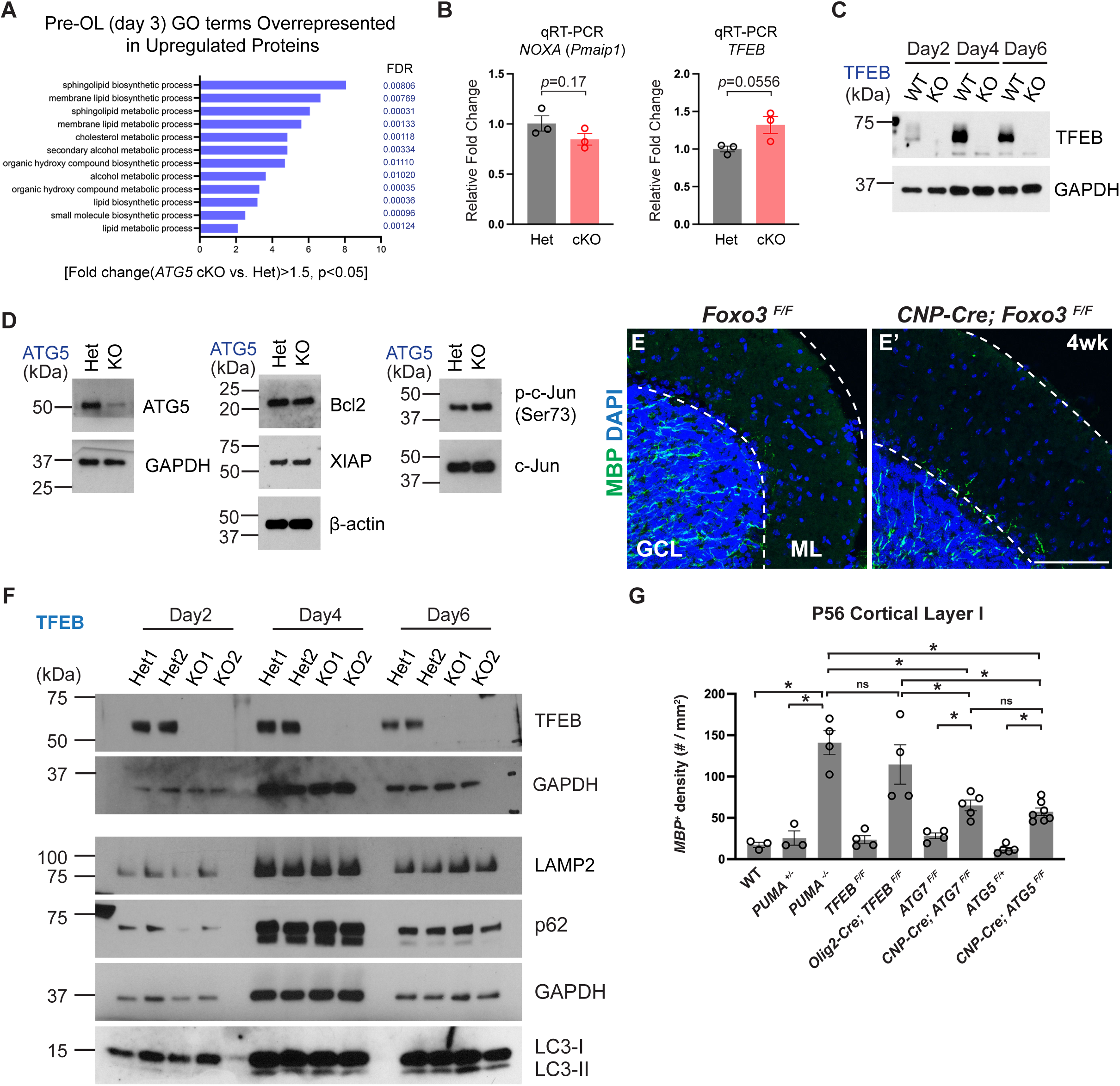
Biochemical characterization of *TFEB*- and *ATG5*-deficient oligodendrocytes and analysis of excessive oligodendrocytes in autophagy and apoptosis mutant mice. **(A)** Gene ontology (GO) term analysis of overrepresented biological processes in *ATG5* cKO pre-OLs as compared to the ones in *ATG5* Het. **(B)** Quantitative RT-PCR analysis of *NOXA*(*Pmaip1*) and *TFEB* mRNA expression in *ATG5* cKO pre-OLs as compared control cells. **(C)** Western blot analysis of *TFEB^F/F^* (WT) and *Olig2-Cre; TFEB^F/F^* (KO) oligodendrocytes during *in vitro* differentiation, showing that TFEB protein is completely abolished in *TFEB* KO oligodendrocytes. **(D)** Western blot analysis of Bcl2, XIAP, c-Jun, and phosphorylated c-Jun (Ser73) in *ATG5* Het and *ATG5* KO pre-OLs, showing that their protein levels remain unchanged in the absence of ATG5. **(E and E’)** Representative confocal micrographs of *Foxo3^F/F^* (E) and *CNP-Cre; Foxo3^F/F^* cerebellum (E’) immunostained by antibodies raised against MBP (green) and counterstained by DAPI (blue), showing that *Foxo3* deletion in oligodendrocyte lineage cells doesn’t result in ectopic myelination in the cerebellar molecular layer. GCL, granule cell layer. ML, molecular layer. **(F)** Western blot analysis of Lamp2, p62, LC3-I and II in *TFEB* Het (*Olig2-Cre; TFEB^F/+^*) and *TFEB* KO (*Olig2-Cre; TFEB^F/F^*) oligodendrocytes during *in vitro* differentiation. *TFEB* KO OLs exhibited similar levels of Lamp2, p62, and LC3I/II as compared to *TFEB* Het OLs throughout *in vitro* differentiation. **(G)** Comparison of *MBP^+^* cell density in cortical layer I between P56 WT, *PUMA^+/-^*, *PUMA^−/−^*, *TFEB^F/F^*, *Olig2-Cre; TFEB^F/F^* (Sun et al., 2018), *ATG7^F/F^*, *CNP-Cre;ATG7^F/F^*, *ATG5^F/+^*, and *CNP-Cre; ATG5^F/F^* (this paper, see Figure 2), showing that *PUMA^−/−^* mutants and *Olig2-Cre; TFEB^F/F^* mutants harbored similar numbers of *MBP^+^* OLs in cortical layer I, however both mouse lines exhibited significantly increased numbers of *MBP^+^* OLs as compared to *CNP-Cre;ATG7^F/F^* and *CNP-Cre; ATG5^F/F^* mutants. Error bars represent SEM. Scale bar: 100 μm in (E’) for (E) and (E’). \**p*<0.05.

**Figure S6, related to Figure 6.**
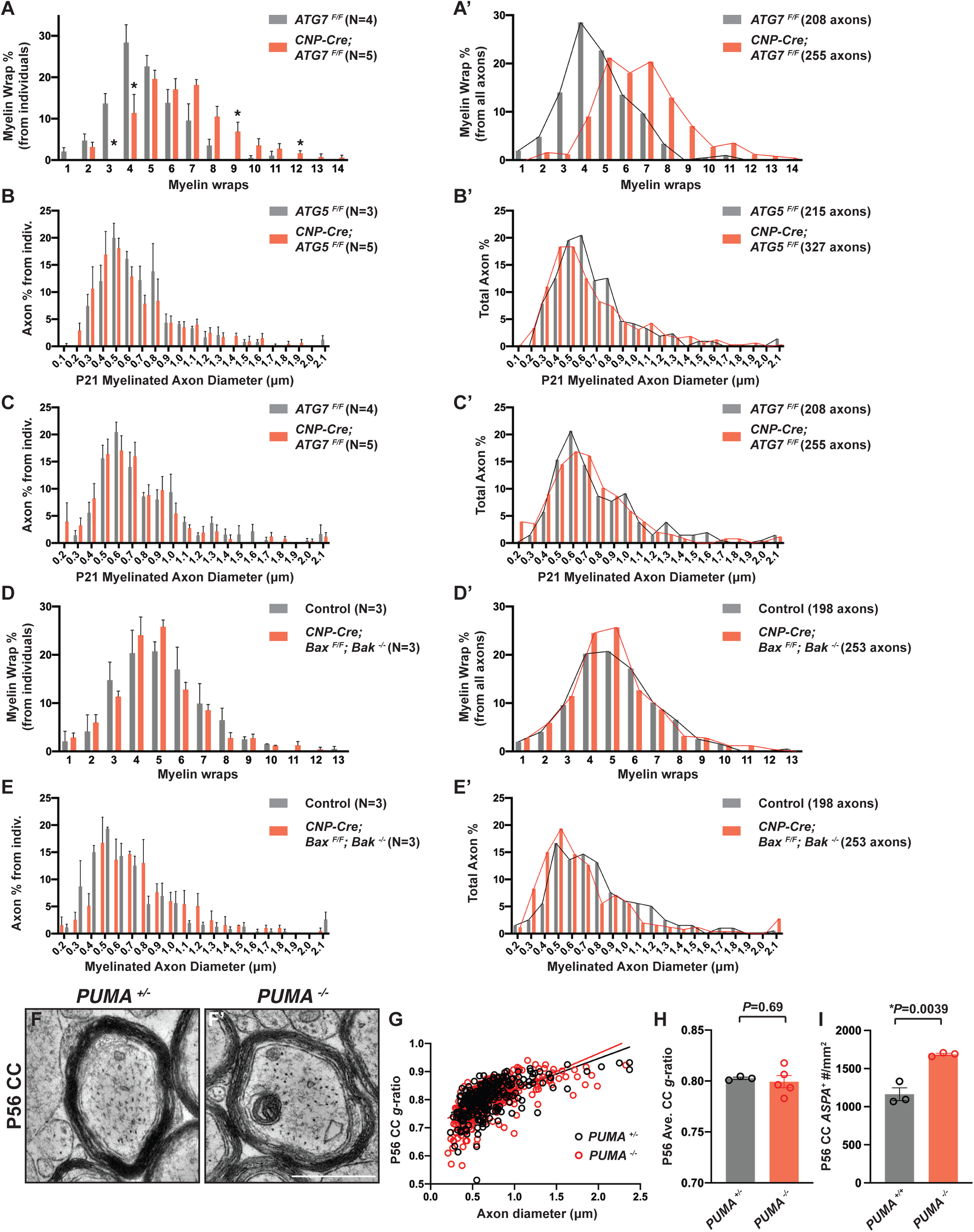
Characterization of axon diameter and myelin sheath number in the *CNP-Cre; ATG7^F/F^*, *CNP-Cre; ATG5^F/F^, CNP-Cre; Bax^F/F^; Bak^−/−^*, and *PUMA^−/−^* mutant mice. **(A and A’)** Quantification of averaged percentage (A) and total percentage (A’) of myelinated axons with different myelin wraps at P21 between *ATG7^F/F^* and *CNP-Cre; ATG7^F/F^* mice. Similar to *CNP-Cre; ATG5^F/F^* mutants (Figures 6J and K), *CNP-Cre; ATG7^F/F^* mutants exhibited a shifted distribution of myelin wrap numbers, showing that decreased *g*-ratios in the *CNP-Cre; ATG7^F/F^* mutants were due to increased myelin wrap numbers. **(B-C’)** Frequency distribution of axons with different diameters in P21 *ATG5^F/F^* and *CNP-Cre; ATG5^F/F^* corpus callosum (B-B’) and in P21 *ATG7^F/F^* and *CNP-Cre; ATG7^F/F^* corpus callosum (C-C’). These results together showed that axon diameter distribution remains unchanged when autophagy is disrupted in oligodendrocytes. **(D-E’)** Frequency distribution of axons with different myelin wrap numbers (D and D’) and frequency distribution of axons with different diameters (E and E’) in the corpus callosum from P21 *CNP-Cre; Bax^F/F^; Bak^−/−^* mutants and littermate controls. These results showed that genetic elimination of pre-OL apoptosis does not change myelin wrap distribution nor alter axon diameter distribution in the corpus callosum. **(F and F’)** Representative TEM micrographs of P56 *PUMA^+/-^* (F) and *PUMA^−/−^* (F’) corpus callosum axons. **(G and H)** Quantification of averaged *g*-ratios of myelinated axons (H), and as a function of axon diameter (G), in P56 *PUMA^−/−^* mutants (red) as compared to littermate controls (black). n≥200 axons from at least 3 animals per genotype in G, and n≥3 animals for each genotype in H. **(I)** Quantification of *ASPA^+^* mature oligodendrocyte numbers in the corpus callosum of P56 *PUMA^+/+^* and *PUMA^−/−^* mice. Error bars represent SEM. Scale bars: 0.5 μm in (F’) for (F) and (F’). \**p*<0.05.

## EXPERIMENTAL MODEL AND SUBJECT DETAILS

### Animals

All experiments related to animals followed protocols approved by the Institutional Animal Care and Use Committee (IACUC) of University of Texas Southwestern Medical Center and University of Texas Health Science Center in accordance with NIH guidelines. Mice were housed on normal light-dark cycles (12:12) with food and water *ad libitum*. The day of birth in this study was designated postnatal day 0 (P0) and all the developmental stages were specified in figures and figure legends. Both male and female mice were used for the experiments and analyses. Transgenic animals include the *ATG7^Flox^* (*ATG7^F^*, RIKEN RBRC02759), *ATG5^Flox^* (*ATG5^F^*, RIKEN RBRC02975), *Olig2-Cre* (Jax#025567), *CNP-Cre* (Lappe-Siefke et al., 2003), *PDGFRα-Cre^ERT2^* (Jax#018280), *TFEB^Flox^* (*TFEB^F^*) (Sun *et al*., 2018), *PUMA^−/−^* (Sun *et al*., 2018)*, Foxo3^Flox^* (*Foxo3^F^*, Jax#024668), and *Bax^Flox^; Bak^−^* (*Bax^F^; Bak^−^*, Jax#006329).

### Immunopanning purification of mouse oligodendrocyte precursor cells (OPCs), *in vitro* differentiation, and Bafilomycin A1 treatment

The immunopanning purification procedure was following the protocol previous described (Sun *et al*., 2018). In brief, postnatal day 6 to day 10 pups were quickly decapitated, and the brains were dissected out and diced into ∼1mm^3^ chunks. The tissues were immediately subjected to digestion with a buffer containing 1× Earle’s Balanced Salt Solution (EBSS 10×; Sigma-Aldrich, Cat#E7501) supplemented with 1 mM MgSO_4_, 0.46% glucose, 2 mM EGTA, 26 mM NaHCO3, 20 units/ml of papain (Worthington Biochemical, Cat#LS003126), and 250 units/ml of DNase I (Worthington Biochemical, Cat#LS002007) under 5% CO_2_/95% O_2_ gas flow at 34°C for 90 minutes. The tissues were then triturated with 1 ml pipettes to form a single-cell suspension. The single-cell suspension was incubated on three BSL-1-coated petri dishes (Vector Laboratories, Cat#L1100) for 10 minutes each to get rid of endothelial cells and microglia. Finally, the suspension was incubated on a rat anti-mouse CD140A (BD Biosciences, Cat#558774) coated petri dish for harvesting OPCs.

The purified OPCs were plated in the proliferation medium containing DMEM-SATO Base Growth Medium (Emery and Dugas, 2013) supplemented with 4.2 mg/ml forskolin (Sigma-Aldrich, Cat#F6886), 10 ng/ml PDGF (Peprotech, Cat#100-13A), 10 ng/ml CNTF (Peprotech, Cat#450-02), and 1 ng/ml neurotrophin-3 (NT-3; Peprotech, Cat#450-03) in a 37°C 10% CO_2_ humidified incubator. To differentiate OPCs *in vitro*, the proliferation medium was replaced with the differentiation medium containing DMEM-SATO Base Growth Medium supplemented with 4.2 mg/ml forskolin, 10 ng/ml CNTF, and 40 ng/ml thyroid hormone (T3; Sigma-Aldrich, Cat#T6397). Half of the culture medium was replaced with fresh medium every 2 days.

To treat cultured oligodendrocytes with bafilomycin A1 (BafA1), 50 nM BafA1 (Sigma-Aldrich, Cat#196000) diluted in DMSO or the same amount of DMSO (vehicle) was added into cell culture medium and cells were incubated at 37°C for 2-3 hours.

### Tamoxifen injections

Tamoxifen was dissolved in sunflower oil at the concentration of 10mg/ml and was injected at postnatal day 4 intraperitoneally with the 100mg/kg dosage.

### *In situ* hybridization and quantification

*In situ* hybridization was performed on fresh frozen sagittal brain sections with 10-μm thickness using a RNAscope Fluorescent Multiplex Reagent Kit (ACDbio, Cat#320850). *MBP in situ* probes (Cat#451491 and Cat#451491-C2), *PDGFRα* probes (Cat#480661-C2) and *ASPA* probes (Cat#425891) were used per manufacturer’s instructions. Fluorescent images were taken using a Keyence BZ-810 all-in-one fluorescence microscope. For the quantification of ectopic oligodendrocytes in the cerebellar molecular layer, the entire cerebellum of each brain section was captured using the automated image-stitching function of Keyence BZ-810. The stitched images were analyzed by the ImageJ software. Briefly, the area of cerebellar molecular layer was selected by the “Freehand Selections” tool followed by the area measurement. The number of *MBP*^+^ cells was counted, and the density was calculated afterwards. *MBP^+^* cell density in the cerebellar molecular layer from at least two brain sections per animal were quantified and averaged, and at least 3 animals per genotype were analyzed.

### Immunohistochemistry

For mouse brain staining, mice were euthanized with CO_2_ and immediately perfused with chilled PBS and 4% PFA in PBS solutions. The brains were post-fixed in 4% PFA/PBS solution for 3 hours at 4°C and then cryopreserved with 30% sucrose in PBS at 4°C for 24 hours. Sagittal brain sections with 18 micron thickness were used for immunostaining. The sections were incubated with 10% normal goat serum (Jackson ImmunoResearch, Cat# 005-000-121) or 10% normal donkey serum (Jackson ImmunoResearch, Cat#017-000-121) in PBS supplemented with 0.1% Triton X-100 (PBST; Sigma-Aldrich, Cat#T8787) for 10-15 minutes at room temperature. The sections were then incubated with the diluted primary antibodies in the staining solution (PBST supplemented with 2% normal goat or donkey serum) at 4°C overnight. The brain sections were rinsed 20 minutes for 4 times in PBST at room temperature and incubated with appropriate AlexaFluor 488- or 594-conjugated secondary antibodies (Thermo Fisher Scientific, 1:1000) and DAPI (Thermo Fisher Scientific, 5 mg/mL, Cat#D1306) for 1 hour at room temperature. The sections were rinsed 20 minutes for 4 times in PBST at room temperature and mounted with coverslips in the VectaShield Hardset Antifade mounting medium (Vector Laboratories, Cat#H-1400-10).

Oligodendrocytes were cultured on Poly-D-lysine (Sigma-Aldrich, Cat# P6407) coated 12-mm plastic coverslips at the density of 10,000 cells/coverslip. Cells were rinsed by PBS and fixed with 4% PFA for 10-15 minutes and rinsed twice with PBS. The cells were then permeabilized and blocked with 10% normal donkey serum in PBST for 10-15 minutes. The cells were incubated with the diluted primary antibodies in PBS supplemented 2% normal donkey serum solution overnight. The cells were washed with PBS 20 minutes for 4 times and incubated AlexaFluor 488- or 594-conjugated secondary antibodies (Thermo Fisher Scientific, 1:1000) and DAPI (Thermo Fisher Scientific, 5 mg/mL) for 1 hour at room temperature or with HCS CellMask Blue Stain (Thermo Fisher Scientific, 1:1000, Cat#H32720) for 1 hour. After 4 times of PBS wash, cells were mounted in the VectaShield Hardset mounting medium. Images were taken by a Keyence all-in-one fluorescence microscope (BZ-X810) or a Zeiss LSM700 inverted confocal microscope.

Primary antibodies used in this study include: rat anti-MBP (Abcam, 1:100 for tissue staining and 1:500 for cell staining, Cat#ab7349), rabbit anti-cleaved Caspase-3 (Asp175) (Cell Signaling Technology, 1:200 for tissue staining and 1:500 for cell staining, Cat#9661S), rabbit anti-PDGFRα (Santa Cruz Biotechnology, 1:200 for tissue staining, Cat#sc-338), and mouse anti-galactocerebroside (Galc) hybridoma (1:50 for cell staining) (Emery et al., 2009).

### Live-cell imaging

OPCs were seeded into PDL-coated 24-well plates at the density of 5,000 cells/well. Cells were maintained in the proliferation medium for 2 days before subjecting to the differentiation medium. For Annexin V imaging, cells were incubated with the differentiation medium supplemented with Alexa 594-conjugated Annexin V (Thermo Fisher Scientific, 1:500, Cat# A13203) or with IncuCyte Annexin V red dye for apoptosis (Saratorius, 1:200, Cat#4641). The plates were placed into IncuCuyte live cell imaging system (Saratorius) and imaged continuously for 5 days with 2-hour intervals at 37°C and 10% CO_2_. Half of the differentiation medium was replaced with fresh differentiation medium supplemented with the indicated dye every 2 days. The Annexin V^+^ area was quantified by the IncuCyte software provided by the vendor. For propidium iodide (PI) staining, the cells were incubated with the differentiation medium for 2 days before the live-cell imaging. At the end of differentiation day 2, half of the medium was replaced with fresh differentiation medium supplemented with PI ((Thermo Fisher Scientific, 1:2000, P1304MP). The plates were placed into IncuCyte live cell imaging system and imaged continuously for 3 days with a 2-hour frame. PI^+^ cell numbers were quantified by the IncuCyte software. At the end of living imaging, Calcein AM (Thermo Fisher Scientific, 1:1000, Cat#C3100MP) or SYTO13 green dye (Thermo Fisher Scientific, 1:10000, Cat# S7575) was added into the medium to label live cells.

### Western immunoblotting and quantification of protein expression

Oligodendrocytes cultured on 6-well plates (100,000 OPCs per plate as the plating density) or 12-well plates (50,000 OPCs per plate as the plating density) were rinsed with chilled PBS and lysed in RIPA Buffer (Thermo Fisher Scientific, Cat#89900) supplemented with 1× cOmplete protease inhibitor cocktail (Sigma-Aldrich, Cat#5892791001) and 1× PhosSTOP phosphatase inhibitors (Sigma-Aldrich, Cat# 4906845001) for 1 hour on a rocker at 4°C. Cell lysates were collected and centrifuged for 10 minutes at 14,000 rpm. The supernatants were collected for use. Protein concentration was determined by a BCA assay (Thermo Fisher Scientific, Cat#23225) and the samples were denatured with 4×LDS sample buffer (Thermo Fisher Scientific, Cat# NP0007) supplemented 10% 2-Mercaptoethanol (Sigma-Aldrich, Cat#M6250) for 10 minutes at 95°C. 1-5 μg total proteins were loaded into stain-free 4-15% or 4%-20% gradient precast polyacrylamide gel (Bio-Rad, Cat# 4568086 or 4568096) for separation. The total protein images were captured by Bio-Rad ChemiDoc imaging system with the stain-free gel imaging protocol. Then proteins were then transferred to 0.45 μM PVDF membranes (Thermo Fisher Scientific, Cat# 88518) at 100V for 1 hour on ice. Blots were blocked with 5% non-fat milk in TBS buffer with 0.1% Tween 20 (TBST) at room-temperature for 1 hour and then incubated with primary antibodies diluted in TBST with 3% BSA overnight at 4°C on a rocker. Blots were washed with TBST for 3X10mintes and incubated with HRP conjugated secondary antibodies in 5% non-fat milk at room-temperature for 1 hour. Blots were washed with TBST for 6 × 5 minutes. Blots were developed with Amersham ECLprime western blotting detection reagent (Cytiva, Cat# RPN2232) for 3 minutes and imaged with X-Ray films. For protein quantification, films were scanned, and quantified with ImageJ and normalized to total proteins, GAPDH or β-actin indicated in the figure legends.

Primary antibodies used include: rabbit anti-TFEB (Bthyl Laboratories, 1:2000, Cat# A303-673A), mouse anti-β-actin (Santa Cruz Biotechnology, 1:500, Cat#sc-47778), rat anti-MBP (Abcam, 1:500, Cat#ab7349), rabbit anti-Beclin1(Cell Signaling Technology, 1:1000, Cat#3495S), rabbit anti-ATG5 (Cell Signaling Technology, 1:1000, Cat#12994S), rabbit anti-ATG7(Cell Signaling Technology, 1:1000, Cat#8558S), rabbit anti-PUMA (Cell Signaling Technology, 1:1000, Cat#24633S), rabbit anti-p62 (Cell Signaling Technology, 1:1000, Cat# 5114S), rabbit anti-Bak (Cell Signaling Technology, 1:1000, Cat# 12105T), rabbit anti-Bax (Cell Signaling Technology, 1:1000, Cat# 2772T), rabbit anti-XIAP (Cell Signaling Technology, 1:200, Cat# 14334S), mouse anti-Bcl2 (Santa Cruz Biotechnology, 1:200, Cat# C2720), mouse anti-WIPI2 (Bio-Rad, 1:1000, Cat# mca5780GA), mouse anti-LC3 (MBL, 1:500, Cat# M186-3), rabbit anti-c-Jun (Cell Signaling Technology, 1:1000, Cat# 9165T), rabbit anti-phospho-c-Jun (Ser73) (Cell Signaling Technology, 1:1000, Cat# 3270T), and mouse anti-GAPDH (Santa Cruz Biotechnology, 1:500, Cat#sc-32233).

### EdU pulse labeling

EdU (5-ethynyl-2’-deoxyuridine) was dissolved in DMSO at a concentration of 10 μg/μl as the stock solution. EdU was administered into P13 *CNP-Cre; ATG7^F/F^* and littermate control animals to pulse-label OPCs via intraperitoneal injections at a dose of 50 μg/g body weight. Mice were perfused with the chilled 4% PFA/PBS solution 24 hours post-injection, and brains were collected and cryopreserved. sagittal brain sections with 16-μm thickness were incubated with the primary antibody solution (rat anti-CD140a, Thermo Fisher Scientific, 1:100, Cat#14140182) at 4°C overnight followed by the incubation with Alexa Fluor 488 secondary antibodies (Thermo Fisher Scientific, 1:1000). The Click-iT® Plus reaction cocktail for EdU detection was then made and applied as described by vendor’s protocol (Thermo Fisher Scientific, Cat#C10639). Fluorescent images were taken using a Keyence BZ-810 all-in-one fluorescence microscope, and the cortical region of each brain section was captured using the automated image stitching

### Transmission electron microscopy and quantification of autophagosomes and lysosomes

Transmission electron microscopy (TEM) of corpus callosum was performed as previously described (Sun *et al*., 2018). For TEM on cultured cells, oligodendrocytes growing on plastic coverslips (Nunc Thermanox Plastic Coverslips, Cat# 174950 Lot#1023669) were fixed with 2.5% (v/v) glutaraldehyde in 0.1M sodium cacodylate buffer with 2mM CaCl_2_ at room temperature for 5 min. The samples were then submitted to the UT Southwestern electron microscopy core for further processing. Briefly, after five rinses in 0.1 M sodium cacodylate buffer, the samples were post-fixed in 1% osmium tetroxide plus 0.8 % K_3_[Fe(CN_6_)] in 0.1 M sodium cacodylate buffer for 1 hour at room temperature. Cells were rinsed with distilled water and *en bloc* stained with 2% aqueous uranyl acetate for 1 hour. After five rinses with distilled water, specimens were dehydrated with increasing concentration of ethanol, infiltrated with Embed-812 resin. Beem capsules were overfilled with resin and coverslips were placed on top, cell side down, and polymerized in a 60°C oven overnight. Epoxy discs were removed by placing coverslips in liquid nitrogen. Beem capsule blocks were sectioned with a diamond knife (Diatome) on a Leica Ultracut UCT (7) ultramicrotome (Leica Microsystems) and collected onto copper grids. The sections were post stained with 2% uranyl acetate in water and lead citrate. Images were acquired on a JEOL 1400+ transmission electron microscope (FEI) equipped with a LaB6 source using a voltage of 120 kV and an AMT camera system.

Imaging was performed at low magnifications to image the whole cell. The low-magnification images were used to calculate whole cell area and nuclear area by tracing their outlines using Fiji. High-magnification images of each cell were taken to quantify autophagosomes and lysosomes. Autophagosomes and autophagosome-like vesicles were identified by their stereotypical double-membrane enclosed structure with various subcellular contents. To prevent inclusion of small vesicles that are likely not autophagosomes in the quantification, all structures above 5000 nm^2^ area were taken for quantification (Smith et al., 2013). Elongated structures were not classified as autophagosomes because these structures could be swollen endoplasmic reticulum (ER). Lysosomes were determined by their stereotypical electron-dense labeling. Individual autophagosomes and lysosomes were counted and manually outlined with the Fiji software to measure their areas. All the quantified organelles were then used to plot graphs with Prism9 software.

### Quantitative mass spectrometry

Oligodendrocytes (∼6,000,000) in two 15-cm dishes were briefly rinsed by DPBS and lysed in 500 μl RIPA buffer supplemented with 1× cOmplete protease inhibitor cocktail and 1× PhosSTOP phosphatase inhibitors. Protein concentration was measured by a BCA assay, and 500 μg proteins were used for quantitative mass spectrometry (MS). The samples were processed at the UT Southwestern Proteomics Core. Briefly, samples were then reduced with tris (2-carboxyethyl) phosphine (TCEP), alkylated with iodoacetamide in the dark, and digested overnight with trypsin at 37°C using an S-Trap (Protifi). Following digestion, the peptide eluate was dried and reconstituted in 100 mM triethylammonium bicarbonate (TEAB) buffer. The samples were labelled with tandem mass tag (TMT) reagent, quenched with 5% hydroxylamine, and combined. The reverse-phase fractionation spin columns (Thermo Fisher Scientific, Cat# 84868) were used according to the manufacturer’s directions to fractionate each sample into 8 fractions. The fractions were dried in a SpeedVac and reconstituted in a 2% acetonitrile, 0.1% trifluoroacetic acid (TFA) buffer. Fractions were injected onto an Orbitrap Fusion Lumos mass spectrometer coupled to an Ultimate 3000 RSLC-Nano liquid chromatography system. Samples were injected onto a 75 μm i.d., 75-cm long EasySpray column (Thermo Fisher Scientifc) and eluted with a gradient 0-28% buffer B over 180 minutes. Buffer A contained 2% (v/v) acetonitrile (ACN) and 0.1% formic acid in water, and buffer B contained 80% (v/v) ACN, 10% (v/v) trifluoroethanol, and 0.1% formic acid in water. The mass spectrometer operated in positive ion mode with a source voltage of 2.0 kV and an ion transfer tube temperature of 275°C. MS scans were acquired at the 120,000 resolution in the Orbitrap and top speed mode was used for SPS-MS3 analysis with a cycle time of 2.5 seconds. MS2 was performed with CID with a collision energy of 35%. The top 10 fragments were selected for MS3 fragmentation using HCD, with a collision energy of 55%. Dynamic exclusion was set for 25 seconds after an ion was selected for fragmentation.

Raw MS data files were analyzed using Proteome Discoverer v2.4 (Thermo Fisher Scientific), with peptide identification performed using Sequest HT searching against the mouse protein database from UniProt. Fragment and precursor tolerances of 10 ppm and 0.6 Da were specified, and three missed cleavages were allowed. Carbamidomethylation of cysteine and TMT labelling of N-terminals and lysine side chains were set as a fixed modification, with oxidation of methionine set as a variable modification. The false-discovery rate (FDR) cutoff was 1% for all peptides.

### RNA extraction, reverse-transcription and quantitative real-time PCR

Oligodendrocytes (∼1,000,000) on 10-cm dishes were rinsed with DPBS and the total RNAs were isolated using the RNeasy Micro Kit (Qiagen, Cat#74004). The total RNAs were reverse-transcribed with the RT SuperMix Kit (New England BioLabs, Cat#E3010) according to vendors’ manuals. The cDNAs were used as the template for quantitative real-time PCR in 20 μL reactions using Fast SYBRGreen Master Mix (Thermo Fisher Scientific, Cat# 4385612) in a QuantStudio 3 qPCR machine (Thermo Fisher Scientific). The primer efficiency was validated by plotting the template quantity vs. the Ct value and individual primer pair’s specificity was confirmed with the single-peak melt-curves. Primer sequences for mouse genes include: *TFEB*-F, CCACCCCAGCCATCAACAC, *TFEB*-R, CAGACAGATACTCCCGAACCTT; *Bbc3* (*PUMA*)-F, ACCTCAACGCGCAGTACG, *Bbc3* (*PUMA*)-R, CACCTAGTTGGGCTCCATTT; *Pmaip1*(*NOXA*)-F, GCAGAGCTACCACCTGAGTTC, *Pmaip1*(*NOXA*)-R, CTTTTGCGACTTCCCAGGCA; *GAPDH*-F, AGGTCGGTGTGAACGGATTTG, *GAPDH*-R, TGTAGACCATGTAGTTGAGGTCA. The quantification was performed with the 2^−ΔΔCt^ method and normalized to *GAPDH*.

### Bulk RNA sequencing (RNA-seq) and analysis

Total RNAs from differentiating oligodendrocytes in 6-well plates (Seeding density: 100,000 OPCs per well) were isolated with the RNeasy Micro Kit (Qiagen, Cat#74004). RNA quality was accessed by Bioanalyzer and samples with high RNA integrity number were used for library construction. Library preparation and messenger RNA (mRNA) sequencing was performed by Novogene Corporation Inc. (Sacramento, California). Differential gene expression analysis was performed using the DESeq2 R package (1.20.0). Genes with an adjusted P-value ≤ 0.05 found by DESeq2 were assigned as differentially expressed genes.

### Compound action potentials (CAPs) measurement on the corpus callosum

2-3 months old mice were used to measure compound action potentials (CAPs). The mice were anesthetized with isoflurane and decapitated. The brain was immediately removed the skull and glued stage of slice chamber containing ice chilled artificial cerebrospinal fluid (aCSF) consisting of 125 NaCl, 2.5 KCl, 1.25 NaH_2_PO_4_, 25 Glucose, 25 NaHCO_3_, 1 MgCl_2_, 2 CaCl_2_ (in mM). The brain was sliced coronally with 400 µm thickness using vibratome (Vt1200S, Leica, German), and slice between bregma 1.10 mm to -0.10 mm were collected. The brain sliced were recovered at aCSF prewarmed at 34L while bubbling with carbogen (95% O_2_ and 5% CO_2_) for 30 min. After recovery the slices were incubated in aCSF at room temperature while bubbling with carbogen (95% O_2_ and 5% CO_2_) until recording. CAPs were recorded in recording chamber which was perfused at 2 ml/min flow rate with room temperature aCSF. Bipolar platinum/iridium customized electrode (FHC, USA, ∼200 µm of distance between positive and negative polars) was placed at 0.5 mm away from midline on the left corpus callosum for stimulation. Recording pipette filled with aCSF (3-6 MΩ 1.0 mm away from midline on right corpus callosum. The latencies of CAPs were measured at midline and at 0.5 mm and 1.0 mm away from midline on right corpus callosum. To measure the size of CAPs, CAPs were recorded at 0.5 mm and 1.0 mm away from midline on right corpus callosum from various strength of stimuli (from 0V to 1V) for 200 µs. Every recording was triplicated and averaged. To calculate conduction velocity, the latencies at 0.5 mm and 1.0 mm away from midline on right corpus callosum were used. The distance was divided by latency difference between two sites.

### Quantification and statistical analysis

All statistical analyses except for the RNA-seq and mass spectrometry analyses (see Method Details) were performed suing GraphPad Prism 9.0. Data are show as mean ± SEM. The “n” numbers for each experiment are specified in the text and figure legends. For the comparison between two groups, statistical significance was determined using two-tailed Student’s *t* test. For multiple comparisons, one-way or two-way ANOVA followed by multiple comparisons tests were performed. The criterion for statistical difference was set at *p*<0.05.

